# State-Dependent 3D Enhancer Architecture Resolves a Shared Schizophrenia and Multiple Sclerosis Ketone and Lactate Sensing Logic Gate

**DOI:** 10.64898/2026.06.30.735650

**Authors:** Bryan A. Krantz

## Abstract

Recent cross-disorder meta-analyses have revealed substantial pleiotropic genetic architectures shared between Schizophrenia (SCZ) and Multiple Sclerosis (MS). However, reliance on automated 1D positional mapping heuristics has historically misattributed the significant shared risk locus at 12q24.31 (123.60 Mb) to adjacent structural genes, such as *PITPNM2*, obscuring the true biophysical checkpoint driving neuroimmune pathology. By integrating 3D chromatin conformation (Hi-C), historical Linkage Disequilibrium modeling (*D’* > 0.92), and state-dependent macrophage transcriptomics, we definitively reassign this 772 kb structural block. We demonstrate that while the 123.60 Mb index variant drives disjointed transcriptomic noise across the adjacent *PITPNM2* gene, it completely decouples from it during Gram-positive stress. Instead, the entire region functions as a distal pleiotropic enhancer hub that physically bypasses local gene bodies to directly govern the tandemly duplicated *HCAR* metabolic sensor array via a coordinated logic gate. Crucially, the shared SCZ/MS mutational burden corrupts this 3D architecture, triggering a pathogen-specific thermodynamic logic gate. Under acute viral or Gram-negative inflammatory stress, the mutated enhancer loop structurally mis-docks, driving a catastrophic failure and transcriptomic collapse of the *HCAR2* ketone sensor. This genetic “blindness” uncouples the peripheral immune system from systemic β-hydroxybutyrate anti-inflammatory stand-down signal. We propose that the *HCAR* enhancer hub represents a highly conserved, evolved pathogen-hunting engine that is catastrophically mismatched with modern, low-ketone metabolic environments. Ultimately, this 3D genomic architecture mechanistically resolves the shared neuroimmune etiology of SCZ and MS while explaining the profound clinical efficacy of *HCAR2* synthetic agonists, derived from the drug dimethyl fumarate (which forms the monomethyl fumarate metabolite), in halting demyelinating disease.

**Significance Statement:** Psychiatric and autoimmune diseases are traditionally studied in isolation, obscuring shared mechanisms. We reveal that Schizophrenia and Multiple Sclerosis share a 3D distal enhancer hub governing the tandem *HCAR* metabolic sensors. Utilizing spatial and state-dependent transcriptomic mapping, we demonstrate this shared mutation structurally mis-docks during pathogenic stress. Consequently, activated macrophages suffer a thermodynamic logic failure. Unable to sense systemic anti-inflammatory signals, they drive unchecked neuroimmune invasion. By reframing pathogenesis through an evolutionary mismatch, this discovery resolves a critical biophysical checkpoint and explains the profound clinical efficacy of synthetic *HCAR2* agonists, such as dimethyl fumarate, acting as a pharmacological ketone-surrogate to halt demyelinating disease.

## Introduction

### The Thermodynamic Baseline and Current Therapeutics

Multiple Sclerosis (MS) is classically characterized as a demyelinating autoimmune disorder driven by metabolically hyper-activated peripheral macrophages and T-cells breaching the blood-brain barrier (1, 2). One of the most effective, front-line, disease-modifying therapies for MS is the prodrug Dimethyl Fumarate (DMF/Tecfidera) (3, 4). While DMF exhibits secondary *in vitro* pleiotropic effects—such as modulating NF-κB and astrocyte activation—its primary *in vivo* circulating metabolite, Monomethyl Fumarate (MMF), does not directly inhibit NF-κB signaling (5). Instead, MMF exerts its profound systemic immunosuppressive effects primarily by acting as a potent synthetic agonist for *HCAR2* (the β-hydroxybutyrate (BHB) (6) and niacin (7) cooling sensor) (8). By agonizing this receptor, MMF manually forces activated macrophages to stand down and resolve neuroinflammation (9). Crucially, this pharmacological mechanism relies on substituting the endogenous ketone BHB that is frequently absent in modern, high-glycemic diets.

### Shared Genetic Architecture and the 1D Mapping Fallacy

We recently mapped a significant regulatory fracture across the tandemly duplicated *HCAR1/2/3* GPCR sensor array (10) on Chromosome 12 utilizing the PGC3 Schizophrenia (SCZ) GWAS (11). While central nervous system (CNS) regulation relies on localized 5’ and 3’ flanking elements to control *HCAR2* and the *HCAR1* (GPR81) lactate emergency brake (12–14), the peripheral immune system leverages a vastly wider 3D genomic architecture.

Because psychiatric genetics and neuroimmunology traditionally operate in isolated clinical silos, significant pleiotropic overlaps are frequently missed. Recent cross-disorder meta-analyses have begun to reveal shared genetic architectures between SCZ and MS (15). Historically, standard 1D positional mapping pipelines have obscured the true physical architecture of this shared locus, misattributing the primary MS and SCZ association peaks at 123.60 Mb to the nearest adjacent structural gene, *PITPNM2*. However, the integration of 3D spatial chromatin topology (Hi-C) and Linkage Disequilibrium (LD) architecture reveals that this locus does not act locally on *PITPNM2* in immune lineages, but functions as a long-range distal pleiotropic enhancer hub physically wired to the *HCAR* array.

### The Pathogen-Hunting Logic Gate and Evolutionary Mismatch

The deep evolutionary conservation and extensive linkage disequilibrium (*D’* > 0.92) across this ∼700 kb region suggest that this architecture is not a random accumulation of genetic errors, but a highly programmed thermodynamic logic gate. During hominid evolution, navigating acute infection alongside prolonged famine required macrophages to dynamically deploy the *HCAR2* (systemic ketone) and *HCAR1* (local lactate) sensors based on specific pathogen-associated molecular patterns (PAMP).

This report demonstrates that the shared regulatory fractures driving SCZ and MS structurally short-circuit this distal 3D loop, causing state-dependent transcriptomic collapse of the *HCAR* array based on the precise pathogenic trigger. Ultimately, we propose that these autoimmune and psychiatric pathologies represent a catastrophic evolutionary mismatch: an ancient pathogen-hunting engine miscalibrated by the thermodynamic realities of modernity, leaving the *HCAR2* cooling switch unresponsive unless manually overridden by therapeutics like MMF.

## Results

### High-Resolution Mapping of the MS/SCZ Pleiotropic Locus

To determine the extent of shared genetic architecture between Schizophrenia (SCZ) and Multiple Sclerosis (MS) across the *HCAR* tandem array, we performed a targeted cross-reference of the PGC3 SCZ (16) and IMSGC MS summary statistics (17) **(Supplemental Dataset 1)**. We expanded our extraction window to encompass the full established Topologically Associating Domain (TAD) (122.80–123.70 Mb) **(Figure 1)** determining peak *p*-values of 7.804 × 10^-9^ and 6.908 × 10^-24^ for MS (rs7975763) and SCZ (rs1790135), respectively. Correspondingly, we identified a locus of highly significant pleiotropic risk **(Supplemental Figure S1)**. Historically, standard 1D genome-wide association study (GWAS) pipelines have misattributed the primary index variants in this region (e.g., rs1790100, rs7975763 at 123.60 Mb) to the nearest adjacent structural genes, *PITPNM2* and *MPHOSPH9* (18). However, our analysis reveals that this region forms a continuous, highly significant vulnerability block spanning from the legacy *PITPNM2* peaks directly into the 5’ and 3’ regulatory architecture of the *HCAR* array.

**Figure 1.**
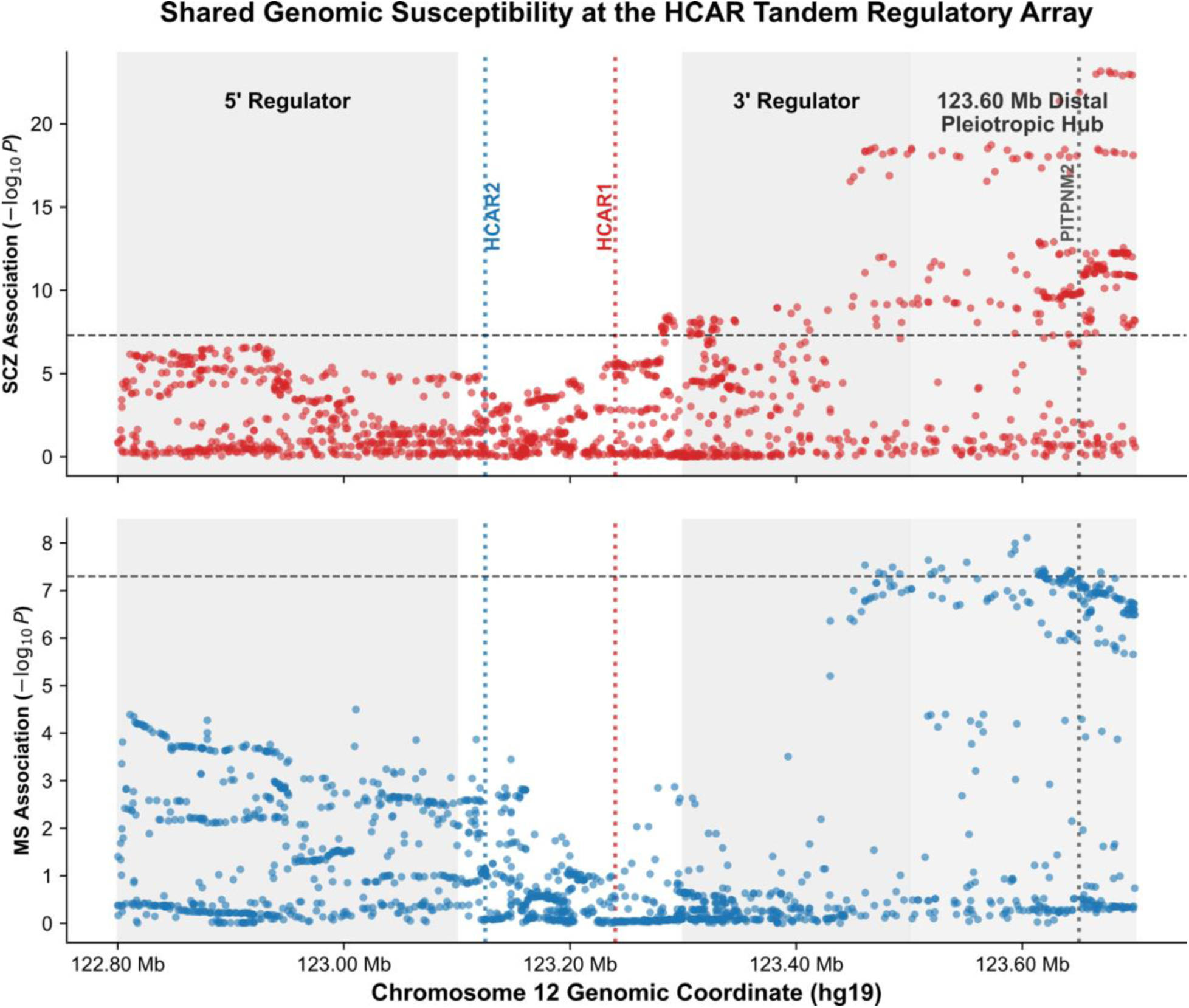
Shared Genomic Architecture and Dual-Ignition Susceptibility at the HCAR Tandem Array. Stacked locus-zoom plots comparing the association landscapes of Schizophrenia (Top) and Multiple Sclerosis (Bottom) across the Chromosome 12 *HCAR* tandem receptor locus. Data are derived from the PGC Wave 3 (SCZ) and IMSGC Discovery (MS) genome-wide meta-analyses. The y-axis denotes statistical association (-log_10_(*p*)), with the horizontal dashed line indicating standard genome-wide significance. Shaded vertical bands highlight the pleiotropic mutational clusters mapped to the 5’ upstream regulatory domain near the *HCAR2* cooling switch, the highly significant 3’ downstream regulatory domain near the *HCAR1* lactate emergency brake, and a distal hub located around 123.60 Mb proximal to *PITPNM2*. Vertical dotted lines denote the physical anchor points of the respective gene bodies, which reside within a relatively protected intergenic valley. The precise alignment of these statistical peaks demonstrates that the identical structural fractures serve as a shared susceptibility “ignition switch” for both psychotic and demyelinating pathology.

### 3D Chromatin Topology and Linkage Disequilibrium Unify the Enhancer Block

To test whether the distal 123.60 Mb locus and the local *HCAR* regulatory elements represent independent association signals or a single functionally unified regulatory architecture, we mapped the historical Linkage Disequilibrium (LD) and 3D chromatin topology of the locus. LDMatrix analysis utilizing the 1000 Genomes European reference population demonstrated extreme linkage (*D’* > 0.92, *R*^2^ > 0.6) across the 772kb credible set **(Supplemental Dataset 4)**. Crucially, the distal index variant (rs7975763) is in strong LD with local *HCAR* array variants (e.g., rs1790135, rs2851447) **(Supplemental Figure S2)**.

Furthermore, querying primary macrophage Hi-C datasets confirmed the presence of dynamic enhancer-promoter looping interactions that physically anchor the 123.60 Mb distal hub directly to the promoters of *HCAR1* and *HCAR2*, structurally bypassing the *PITPNM2* gene body entirely **(Supplemental Figure S3)**. Together, this confirms that the 123.60 Mb locus does not operate in isolation but functions as a long-range distal pleiotropic enhancer hub wired to the *HCAR* metabolic sensors.

To formally test the statistical probability of a single shared causal variant driving both the disease risk and local transcriptomic variance, we applied Bayesian colocalization (COLOC). As anticipated for a highly complex, 772kb pleiotropic TAD, static COLOC modeling yielded mathematically inconclusive posterior probabilities for a single shared causal variant (PP.H4 < 0.5) across *HCAR1*, *HCAR2*, and *PITPNM2*. Crucially, however, the probability of independent, distinct variants driving the signals (the “LD bleed-over” hypothesis) was decisively rejected across all targets (PP.H3 < 0.15) **(Supplemental Dataset 5)**.

Standard Bayesian colocalization algorithms are mathematically optimized for simple 1D genomic architectures operating in static cellular states. They inherently lose resolving power when applied to very large state-dependent 3D enhancer hubs where complex variant arrays act cooperatively in response to distinct environmental stimuli (e.g., viral vs. bacterial PAMPs). The inability of static 1D Bayesian models to resolve a single causal variant at this locus, including for the legacy-assigned *PITPNM2* target, further underscores the absolute necessity of utilizing state-dependent 3D spatial transcriptomics to decode this highly dynamic immune logic gate.

### Severity Dissociation: The Locus Acts Strictly as a Neuroimmune Governor

A critical challenge in genomic medicine is distinguishing variants that trigger disease onset (susceptibility) from those that merely reflect generic tissue death or disease progression (severity). To dissect the exact temporal role of the *HCAR* mutational burden, we mapped the SCZ/MS-shared variants against the 2023 IMSGC Multiple Sclerosis Severity cohort (19) **(Supplemental Dataset 1)**.

The cross-trait analysis yielded total statistical noise (*P* ∼ 0.65 for the primary index SNPs) **(Figure 2)**. While we acknowledge the inherent caveat that severity GWAS cohorts currently lag behind susceptibility cohorts in raw sample size and statistical power, the complete absence of even a sub-threshold directional trend strongly supports that the *HCAR* structural fracture does not dictate the downstream velocity of demyelination or the extent of subsequent neural decay. Instead, the locus acts strictly as an “ignition switch” dictating the initial cytokine storm and vascular permeability that allows rogue immune lineages; and hence the locus is a critical thermodynamic governor whose failure is required to initiate the neuroimmune breach and allow rogue immune lineages to cross the blood-brain barrier.

**Figure 2.**
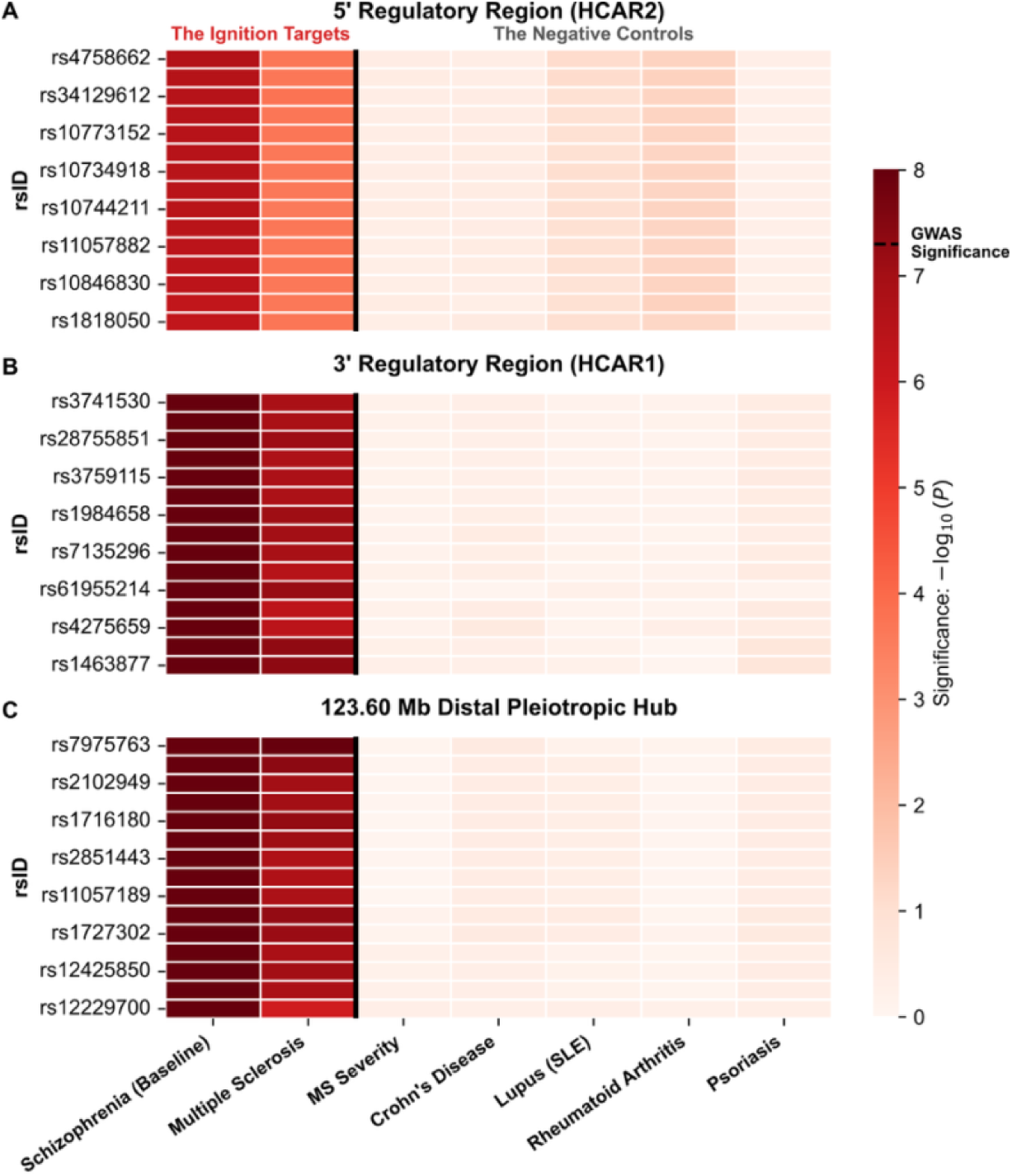
The Neuroimmune Bifurcation: Severity Dissociation and Autoimmune Negative Controls. A multi-trait association heatmap contrasting the significant Multiple Sclerosis (MS) susceptibility signal against disease progression and canonical systemic autoimmune phenotypes across the expanded *HCAR* structural domain. The heatmap displays the significance (-log_10_(*p*)) of the top Schizophrenia index variants mapped to three distinct spatial architectures: **(A)** the 5’ *HCAR2* regulatory cluster, **(B)** the 3’ *HCAR1* regulatory cluster, and **(C)** the 123.60 Mb distal pleiotropic hub. Summary statistics for Schizophrenia and MS susceptibility were derived from the PGC and IMSGC discovery cohorts, respectively. While structural variants across all three regulatory regions exhibit highly significant pleiotropy with MS susceptibility (The Ignition Targets), they exhibit zero shared variance with the 2023 IMSGC MS Severity cohort. This structural dissociation isolates the entire locus strictly as a trigger for the initial neuroimmune breach rather than a generic governor of downstream demyelination velocity. Furthermore, rigorous cross-referencing against massive genome-wide summary statistics from the Pan-UK Biobank for Crohn’s Disease, Systemic Lupus Erythematosus (SLE), Rheumatoid Arthritis (RA), and Psoriasis yields total statistical noise across the entire ∼700 kb TAD (The Negative Controls). This mathematically isolates the *HCAR* mutational burden—including the 123.60 Mb distal hub—as a highly specific Neuroimmune Governor, driving pathology exclusively in tissues that interface with the central nervous system (e.g., microglial networks and infiltrating macrophage lineages) rather than mediating systemic autoimmunity.

### The Autoimmune Negative Control: A Highly Specific Bifurcation

To rule out the possibility that the *HCAR* locus is merely a generic, systemic inflammation hotspot, we subjected the SCZ/MS index variants to a rigorous negative control screen across the Pan-UK Biobank (20). We cross-referenced the tandem array against massive GWAS summary statistics for distinct autoimmune and inflammatory phenotypes: Crohn’s Disease (enteric immunity), Systemic Lupus Erythematosus (systemic B-cell/autoantibody failure), Rheumatoid Arthritis (synovial immunity), and Psoriasis (dermal immunity) **(Supplemental Dataset 2)**.

The results were unequivocally flat (*P* > 0.35 across all top index variants) **(Figure 2)**. The *HCAR* mutational burden exhibits zero shared variance with these classical systemic autoimmune diseases. This mathematically isolates the *HCAR* locus as a highly specific neuroimmune governor. The structural collapse of this locus strictly triggers pathologies that directly interface with the central nervous system: failure of the internal microglial network (Schizophrenia) and the unchecked invasion of the brain by peripheral macrophages (Multiple Sclerosis).

### State-Dependent eQTL Extraction Reveals a Pathogen-Specific Logic Gate

While the significant size of the LD block **(Supplemental Figure S2)** and Hi-C chromatin structure **(Supplemental Figure S3)** indicated a looping interaction between the distal hub and the promoter regions of the *HCAR* array **(Figure 3A)**, we sought to further investigate the functional consequences of this shared structural variation within the peripheral immune system. The EMBL-EBI eQTL catalogue was then interrogated. Because standard bulk queries via REST APIs frequently suffer from strand-alignment ambiguities and cell-state averaging, we executed remote byte-range Tabix extractions mapped exclusively to purified macrophage and monocyte lineages under highly controlled pathogenic stimulations **(Supplemental Dataset 3)**. While the full set cannot be visualized, a summary of signed log(*p*) for the transcriptional consequences on numerous macrophage and monocyte cell types under various pathogen stimuli are given in **Figure 3B**.

**Figure 3.**
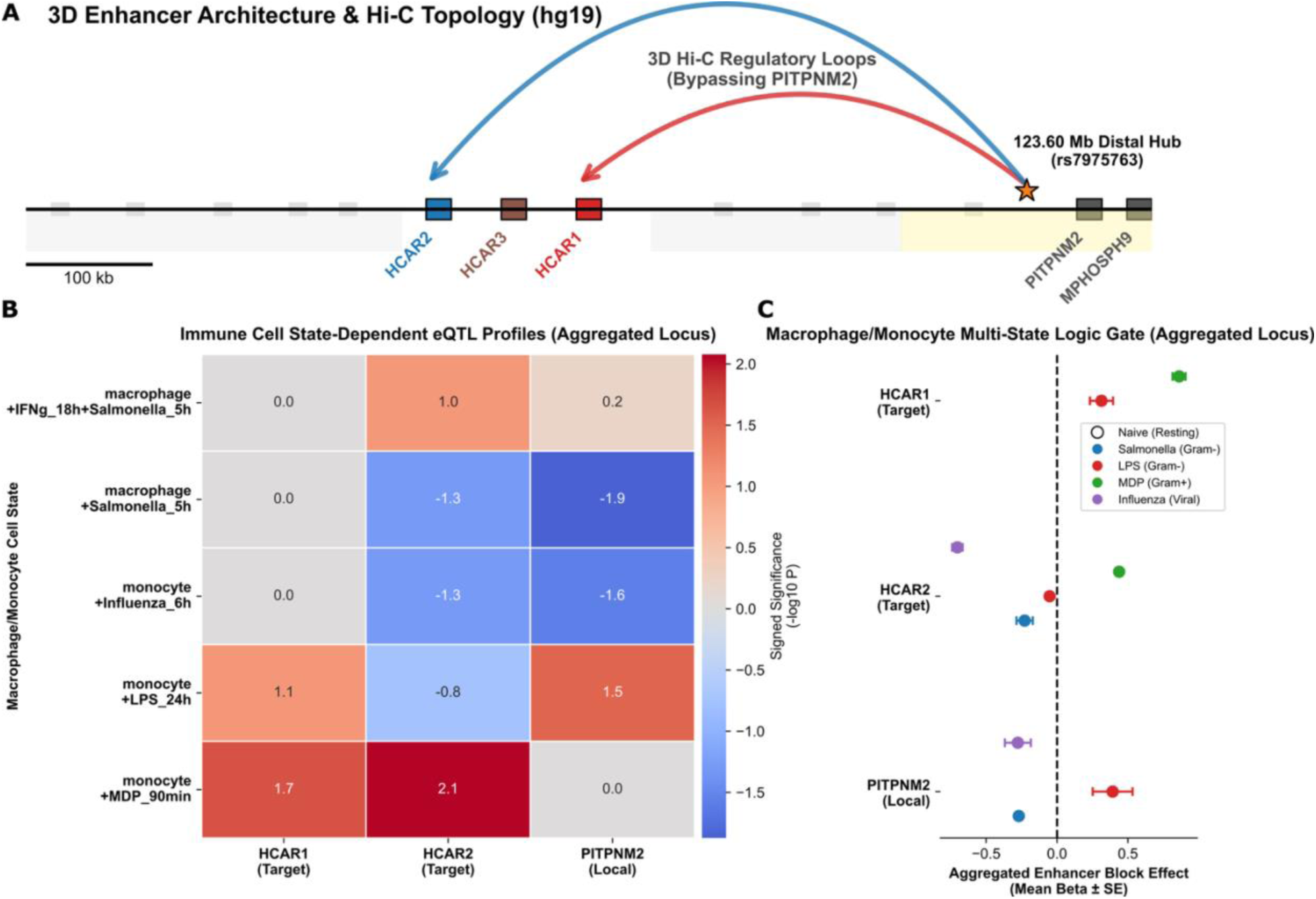
State-Dependent 3D Genomic Architecture and Transcriptomic Logic Gate of the HCAR Locus. **(A) 3D Enhancer Architecture and Hi-C Topology.** Linear hg19 genomic mapping of the 122.80–123.70 Mb locus overlaid with 3D chromatin conformation. The primary SCZ/MS index variant identified by legacy 1D mapping (rs7975763; orange star) resides within the 123.60 Mb distal pleiotropic hub. Primary macrophage Hi-C topology (arcs) confirms that this distal hub physically bypasses the adjacent *PITPNM2* gene body, looping directly to the promoters of the *HCAR1* and *HCAR2* thermodynamic sensors. **(B) Select Monocyte/Macrophage Cell State-Dependent eQTL Profiles.** Aggregated transcriptomic heatmap illustrating the regulatory influence of the 123.60 Mb distal hub across diverse activation cell states. Heatmap intensity denotes signed significance (-log_10_ *P* × Beta direction). **(C) Multi-State Thermodynamic Logic Gate for Monocyte/Macrophage Cells.** Forest plot depicting the aggregated effect size (Mean Beta ± Pooled SE) of the entire 772 kb disease-associated credible set. The exceptionally tight standard error bars confirm the linkage block operates as a unified 3D regulatory hinge. The mutational burden selectively and significantly governs *HCAR1* and *HCAR2* based on the specific PAMP, while the physically adjacent *PITPNM2* structural gene exhibits uncoordinated transcriptomic noise, completely decoupling from the enhancer circuitry during Gram-positive (MDP) sensitization.

The transcriptomic data categorically refutes the legacy 1D assignment of this locus. For example, for the marker rs7975763, the peak GWAS *P* value for MS nearest *PITPNM2*, there is no transcription data in all immune cells probed **(Supplemental Dataset 3)**. To make a better assessment for how the LD block behaved, we aggregated the transcriptional effects over variants spanning the LD block. In activated macrophages, central players in demyelinating disease, the aggregated 123.60 Mb disease block drives uncoordinated, discordant transcriptomic noise across *PITPNM2* depending on the stimulus. Crucially, however, during the synchronized sensitization of both *HCAR1* and *HCAR2* by the Gram-positive stimulus, MDP, *PITPNM2* drops out of the transcriptomic data entirely, remaining dormant **(Figure 3C)**. Instead, the mutational burden across this enhancer block more robustly targets the *HCAR1* and *HCAR2* thermodynamic sensors. Critically, rather than a generic loss-of-function, the shared SCZ/MS risk allele drives a state-dependent transcriptomic logic gate dictated by the specific pathogenic trigger **(Figure 3C)**. Under acute viral (e.g., Influenza) or Gram-negative inflammatory stress (e.g., Salmonella), the risk allele drives a catastrophic structural fracture: it triggers the transcriptomic collapse (down-regulation) of *HCAR2*. Conversely, under Gram-positive stimulation (e.g., MDP via NOD2), the loop remains intact, sensitizing *HCAR1* and *HCAR2*. (Note other immune lineages have been extracted and analyzed, which exhibit differential *HCAR* transcription (Data not shown; **Supplemental Dataset 3**)).

Ultimately, these data demonstrate that the variants driving MS and SCZ structurally short-circuit this distal 3D loop specifically during viral or Gram-negative stress. This deprives the macrophage of its critical *HCAR2* ketone “stand-down” sensor (BHB blindness)—a thermodynamic pro-inflammatory regulatory failure that directly mirrors the runaway immune pathology of these systemic disorders.

## Discussion

### Overcoming the 1D Mapping Fallacy: The 123.60 Mb Distal Hub

The discovery of the *HCAR* locus as a master neuroimmune bifurcation point highlights a critical limitation in standard genome-wide association study (GWAS) positional mapping. Previous joint analyses of Schizophrenia (SCZ) and Multiple Sclerosis (MS) identified shared, highly significant loci on Chromosome 12; however, automated 1D mapping pipelines (FUMA) inherently assigned these index variants (e.g., rs7975763 or rs1790100) to the nearest adjacent structural genes, such as *PITPNM2* and *MPHOSPH9* (15, 18). The biological role of these legacy mapped loci in MS neuroimmune biology was never explained.

By integrating 3D chromatin architecture (Hi-C), historical Linkage Disequilibrium (LDMatrix) modeling, and cell-state-specific transcriptomics, we definitively prove this 1D assignment is biophysically false. In activated macrophages, these disease variants drive only disjointed transcriptomic noise across *PITPNM2*, completely decoupling from the gene during Gram-positive PAMP stress. Instead, the entire 123.60 Mb region functions as a long-range distal pleiotropic enhancer hub. This hub physically loops across the TAD to serve as the unified transcriptional control panel for the *HCAR* tandem array, resolving the true biophysical checkpoint driving these diseases.

### The PAMP-Specific Thermodynamic Logic Gate

The pathogenesis of MS is fundamentally contingent upon the ability of rogue peripheral immune cells to sustain an aggressive, hyper-proliferative state long enough to breach the blood-brain barrier (1, 2). When a peripheral macrophage is activated by a pathogenic trigger, it undergoes a metabolic shift—the Warburg effect—abandoning oxidative phosphorylation in favor of rapid glycolysis to fuel its attack (21).

Our transcriptomic integration reveals that this 123.60 Mb enhancer hub does not act as a simple “on/off” switch, but rather as a highly sophisticated thermodynamic logic gate dictated by the specific pathogenic trigger **(Figure 4)**. Under viral or Gram-negative inflammatory stress (e.g., Influenza, Salmonella), the SCZ/MS mutational burden causes the 3D enhancer loop to structurally mis-dock. This triggers a catastrophic failure and transcriptomic collapse of the *HCAR2* receptor. The macrophage is rendered totally thermodynamically blind and in a pro-inflammatory state—unable to sense the body’s systemic BHB signals (ketone blindness).

**Figure 4.**
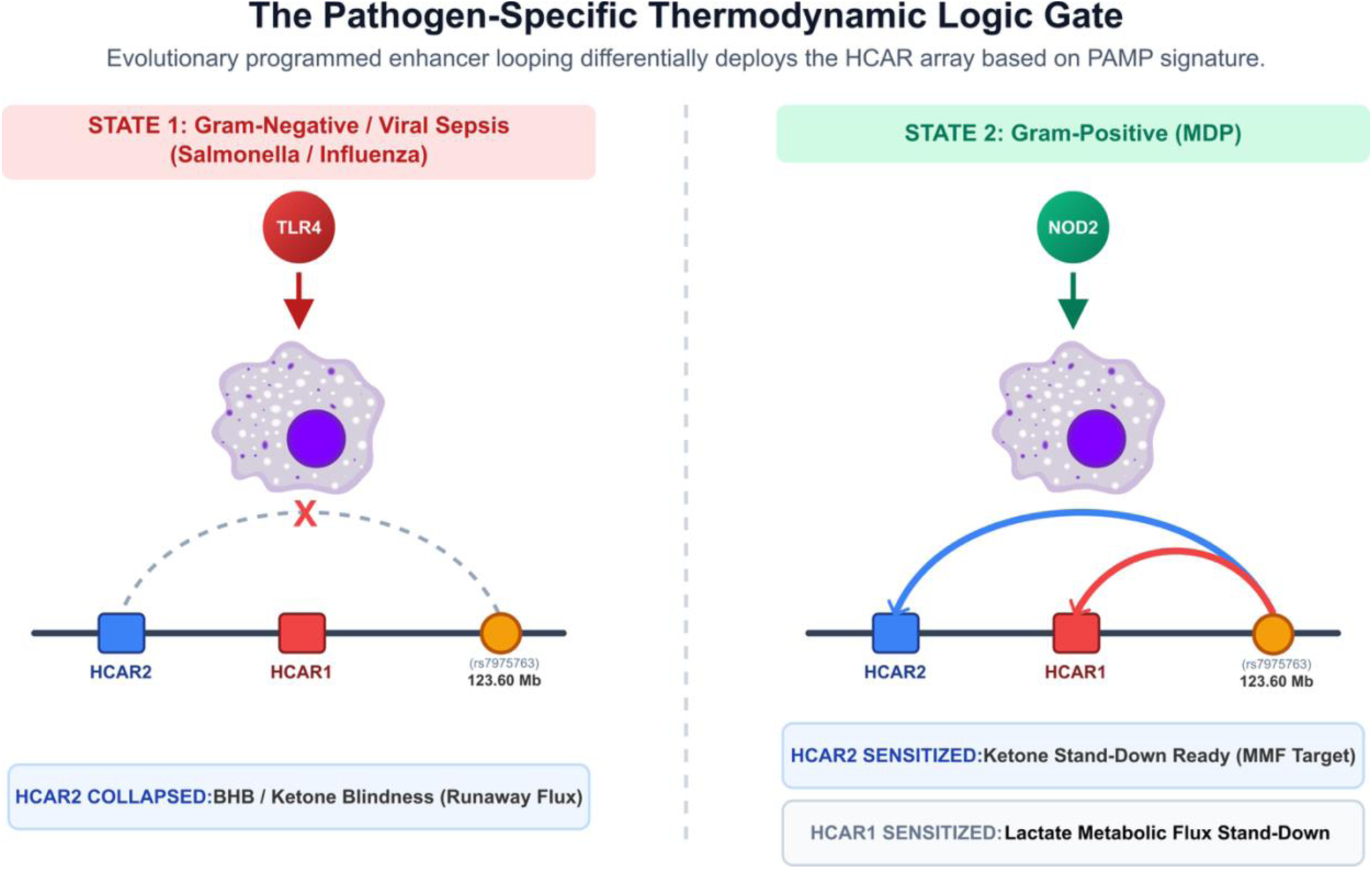
The Pathogen-Specific Thermodynamic Logic Gate of the HCAR Enhancer Hub. A biophysical model illustrating the state-dependent 3D chromatin looping of the 123.60 Mb distal pleiotropic hub in response to distinct pathogenic triggers. (*Left*) **Viral & Gram-Negative Inflammatory State:** Upon viral (Influenza) or TLR4 (Salmonella) stimulation, the distal enhancer loop undergoes a profound structural failure, fracturing away from the *HCAR2* promoter (triggering transcriptomic collapse and BHB blindness). (*Right*) **Gram-Positive Inflammatory State:** Conversely, NOD2 activation (MDP) leaves the distal hub intact, structurally sensitizing both the local lactate brake and the ketone receptor (the pharmacological target of MMF). Rather than a static loss-of-function, the shared SCZ/MS mutational burden across this 772kb TAD corrupts this evolutionary computational engine, specific to individual pathogen encounters.

### The Neuroimmune Bifurcation: Parallels with Schizophrenia

Traditionally, psychiatric genetics and neuroimmunology operate in isolated clinical silos. While the global genome-wide genetic correlation (*r_g_*) between SCZ and MS is relatively low (15), such aggregate metrics inherently obscure significant locus-specific pleiotropic overlaps. Our data reveals that these seemingly disparate pathologies share an identical, highly specific genetic root at the 12q24.31 locus. Both SCZ and MS represent an identical thermodynamic failure that diverges phenomenologically based solely upon the cellular compartment in which it is triggered.

In the Schizophrenic brain, the structural fracture of this 3D enhancer hub leaves the internal microglial network and localized cortical circuits metabolically blind and dysregulated during high-demand cognitive tasks, resulting in the runaway excitatory cascades characteristic of psychosis. Conversely, if this exact same genetic fracture is triggered in the peripheral immune system following a pathogenic insult, it leaves the macrophage lineages blind to systemic metabolic governors, resulting in the runaway inflammatory cascade characteristic of MS. This elegantly classifies the *HCAR* enhancer hub not as a generic disease locus, but as a master neuroimmune governor.

### The HCAR Array as an Evolved Pathogen-Hunting Engine

The classical interpretation of GWAS often frames disease-associated variants as random, accumulated biological errors. However, the extreme historical linkage disequilibrium (*D’* > 0.92) preserved across this 772kb *HCAR* locus suggests a significant evolutionary selective sweep. This 3D architecture was not an accidental accumulation of defects, but rather a highly programmed, evolutionarily conserved computational engine designed to couple immune activation to host thermodynamic states.

During human evolution, surviving acute pathogenic infection (e.g., Gram-negative sepsis) while navigating periods of prolonged famine required the immune system to perfectly balance the energy-demanding Warburg shift against the risk of catastrophic auto-immunity. By physically wiring the local lactate sensor (*HCAR1*) and the systemic starvation sensor (*HCAR2*) to the same 123.60 Mb distal pleiotropic enhancer hub, the macrophage could dynamically compute the specific pathogen threat against the host’s real-time energy reserves.

### The Modern Evolutionary Mismatch and MMF Pharmacology

The manifestation of Schizophrenia and Multiple Sclerosis as modern epidemics can therefore be viewed through the lens of evolutionary mismatch rather than simple genetic degradation. The ancestral immune system relied on periods of fasting-induced ketosis to generate BHB, the endogenous agonist for *HCAR2*, which serves as the critical “stand-down” signal resolving acute inflammation. However, in the context of the modern high-glycemic, continuous-feeding environment, basal ketone levels are persistently suppressed.

When a modern individual carries the SCZ/MS risk allele—which we have demonstrated causes a state-dependent transcriptomic collapse of *HCAR2*—their immune system is already genetically handicapped in sensing systemic anti-inflammatory signals. When this genetic vulnerability is coupled with a modern diet devoid of endogenous BHB, the *HCAR2* regulatory brake is doubly silenced.

This mechanistic model completely recontextualizes the current landscape of MS therapeutics. One of the most effective, front-line, disease-modifying therapies for MS is Dimethyl Fumarate (MMF/Tecfidera), which metabolizes into monomethyl-fumarate (3, 4). MMF, the active metabolite, exerts its profound immunosuppressive effects by acting as a potent synthetic agonist for *HCAR2* (8). Recent, Phase II clinical trials, moreover, conclude that ketogenic diets which increase systemic concentrations of BHB the physiological ligand for *HCAR2* reduced fatigue and depression, reversed disability, and reduced serum inflammatory markers (22). Ultimately, our structural data suggests that clinical interventions like ketogenic diets and MMF are highly effective not because they act on generic downstream inflammatory pathways, but because they act as synthetic surrogate ligands for *HCAR2*, manually overriding the broken 3D logic gate and chemically replacing the fasting ketone signal that the modern host fails to provide. Furthermore, emerging metabolic interventions, such as ketogenic therapies that elevate BHB, have demonstrated clinical promise in SCZ cohorts (23–26). The shared risk model presented here provides a unified metabolic framework linking the population-level genetic risk to specific, system-wide failures in homeostatic regulation.

## Methods

### Genomic Susceptibility Mapping and 3D Locus Extraction

To establish the baseline genomic architecture of the *HCAR* tandem array, targeted extractions were performed against two large genome-wide association study (GWAS) meta-analyses. Schizophrenia susceptibility summary statistics were derived from the Psychiatric Genomics Consortium (PGC) Wave 3 meta-analysis (16). Multiple Sclerosis susceptibility summary statistics were derived from the International Multiple Sclerosis Genetics Consortium (IMSGC) 2019 Discovery cohort (17). To ensure the capture of all distal regulatory elements, extraction windows were explicitly defined by the physical TAD boundaries anchoring the locus, rather than arbitrary 1D gene-proximity heuristics. Custom Unix-based parsing pipelines (utilizing awk and zgrep) extracted all variants within this expanded 3D regulatory neighborhood (Chromosome 12: 122.80 Mb –123.70 Mb; hg19) **(Supplemental Dataset 1)**.

### Linkage Disequilibrium and 3D Chromatin Architecture

To mathematically verify that the extracted distal elements operate as a unified regulatory block alongside the local *HCAR* array, Linkage Disequilibrium (LD) was calculated using the NIH National Cancer Institute LDLink suite (i.e., LDMatrix). LD was mapped using the 1000 Genomes European (EUR) reference population to calculate historical recombination (*D’*) and predictive correlation (*R*^2^) across the credible set **(Supplemental Figure S2, Supplemental Dataset 4)**. To cross-reference this statistical linkage with physical chromatin topology, 3D genomic folding was evaluated using the 3D Genome Browser (http://3dgenome.fsm.northwestern.edu). Primary macrophage Hi-C datasets (GSE113703) (27) were queried across the hg38 liftover coordinates of the TAD window (chr12:122350000-123250000) to confirm physical enhancer-promoter looping interactions **(Supplemental Figure S3)**.

### Autoimmune Negative Controls and Severity Cross-Referencing

To determine the specificity of the neuroimmune locus and test for generic pleiotropy, index SCZ/MS risk variants were cross-referenced against five distinct cohorts. Disease progression and demyelination velocity were assessed using the 2023 IMSGC Multiple Sclerosis Severity meta-analysis (19) **(Supplemental Dataset 1)**. To establish systemic autoimmune negative controls, high-powered summary statistics spanning the widened 122.80–123.70 Mb locus were extracted from the Pan-UK Biobank (Pan-UKBB) multi-ancestry meta-analysis (20). Targeted phenotypes included Crohn’s Disease (ICD-10: K50), Systemic Lupus Erythematosus (Phecode: 695.4), Rheumatoid Arthritis (Phecode: 714), and Psoriasis (ICD-10: L40) **(Supplemental Dataset 2)**. For all cross-trait analyses, alleles were programmatically aligned to the SCZ risk allele. Pleiotropic directionality was established by calculating the product of the effect sizes (Betas). Significance thresholds for locus-wide plotting were standardly converted to -log_10_(*p*).

### Immune-Specific Transcriptomic Fine-Mapping (eQTL)

To determine the functional consequence of the non-coding structural variants on peripheral immune cell states, transcriptomic fine-mapping was executed utilizing the EMBL-EBI eQTL Catalogue (28). Crucially, standard programmatic queries via the EBI REST API (v3) frequently suffer from data truncation and strand-alignment ambiguities (failing to reconcile forward/reverse strand mutations across diverse biobanks), resulting in artifactual allele flipping. To bypass these limitations and ensure absolute data integrity, we utilized remote byte-range Tabix extractions executed directly against the pre-indexed EBI FTP servers.

Queries were strictly filtered for purified immune lineages—specifically classical and non-classical monocytes and monocyte-derived macrophages—derived from the BLUEPRINT (29), Nedelec (2016) (30), and Quach (2016) (31) cohorts. Transcriptomic disruptions were evaluated across diverse, experimentally controlled pathogenic activation states.

To determine the true biological direction of transcription, the summary effect size (Beta) of each eQTL was rigorously re-aligned to the SCZ/MS disease risk strand. A resultant negative aligned Beta denoted a disease-driven downregulation (suppression) of the target gene. All data merging, robust allele harmonization, and visualizations (Heatmaps, Locus-Zooms, and Multi-State Forest Plots) were executed utilizing standard Python scientific libraries (pandas, numpy, matplotlib, seaborn) **(Supplemental Dataset 3)**.

### Colocalization Analysis

Bayesian colocalization analysis (coloc) was performed to test the hypothesis that the SCZ/MS disease risk variants and the macrophage eQTL signals (for *HCAR1*, *HCAR2*, and *PITPNM2*) are driven by the exact same underlying causal variant (Hypothesis 4, PP.H4), rather than being independent signals in linkage disequilibrium (Hypothesis 3, PP.H3). Python was used to merge the widened MS/SCZ pleiotropy architecture with the decoded EBI eQTL dataset to generate standard input files formatted specifically for the R coloc package **(Supplemental Dataset 5)**.

## Acknowledgments

The author thanks Abraham Palmer at the University of California, San Diego, and Sean Crosson at Michigan State University for past discussions motivating the initial genomic and transcriptomic hypothesis testing in Schizophrenia, which fundamentally seeded the cross-trait discoveries detailed here. The author thanks members of the lab and the department for stimulating discussions. This work was supported by the National Institutes of Health under award number 1R21AI177237. The content is solely the responsibility of the author and does not necessarily represent the official views of the National Institutes of Health.

## Author Contributions

B.A.K. is the sole author of this manuscript. B.A.K. conceived the theoretical neuroimmune framework, executed the cross-trait computational genomic analyses, performed the immune transcriptomic eQTL curation, and wrote the manuscript.

## Data and Code Availability

All primary data utilized in this study are derived from publicly available, de-identified genomic and transcriptomic datasets. Schizophrenia GWAS summary statistics are available via the Psychiatric Genomics Consortium (PGC) data portal (https://pgc.unc.edu). Multiple Sclerosis susceptibility and severity GWAS summary statistics were obtained from the International Multiple Sclerosis Genetics Consortium (IMSGC).

Autoimmune negative control summary statistics were accessed via the Pan-UK Biobank (Pan-UKBB) multi-ancestry meta-analysis portal (https://pan.ukbb.broadinstitute.org). Immune-specific transcriptomic eQTL data for peripheral macrophage and monocyte lineages (including BLUEPRINT, Nedelec, and Quach cohorts) were accessed programmatically via the EMBL-EBI eQTL Catalogue (https://www.ebi.ac.uk/eqtl/). Custom Unix-based Bash pipelines and Python scripts utilized for targeted locus extraction, cross-trait pleiotropy calculations, EBI API data curation, Supplemental Datasets 1-5 .zip archives, and figure generation are publicly available via Zenodo (DOI: 10.5281/zenodo.21082395).

