## Supplemental Data for "State-Dependent 3D Enhancer Architecture Resolves a Shared Schizophrenia and Multiple Sclerosis Ketone and Lactate Sensing Logic Gate"

Bryan A. Krantz

Department of Microbial Pathogenesis, School of Dentistry, University of Maryland, Baltimore, 650 W. Baltimore Street, Baltimore, MD 21201, U.S.A.

 (BAK)

**Running title:** Shared HCAR Gates in SCZ and MS

**Keywords:** Multiple Sclerosis, Schizophrenia, Lactate, Beta-Hydroxybutyrate (BHB), Genomics, Transcriptomics, eQTL, HCAR1 (GPR81), HCAR2 (GPR109A), Dimethyl Fumarate, Monomethyl Fumarate, Tecfidera

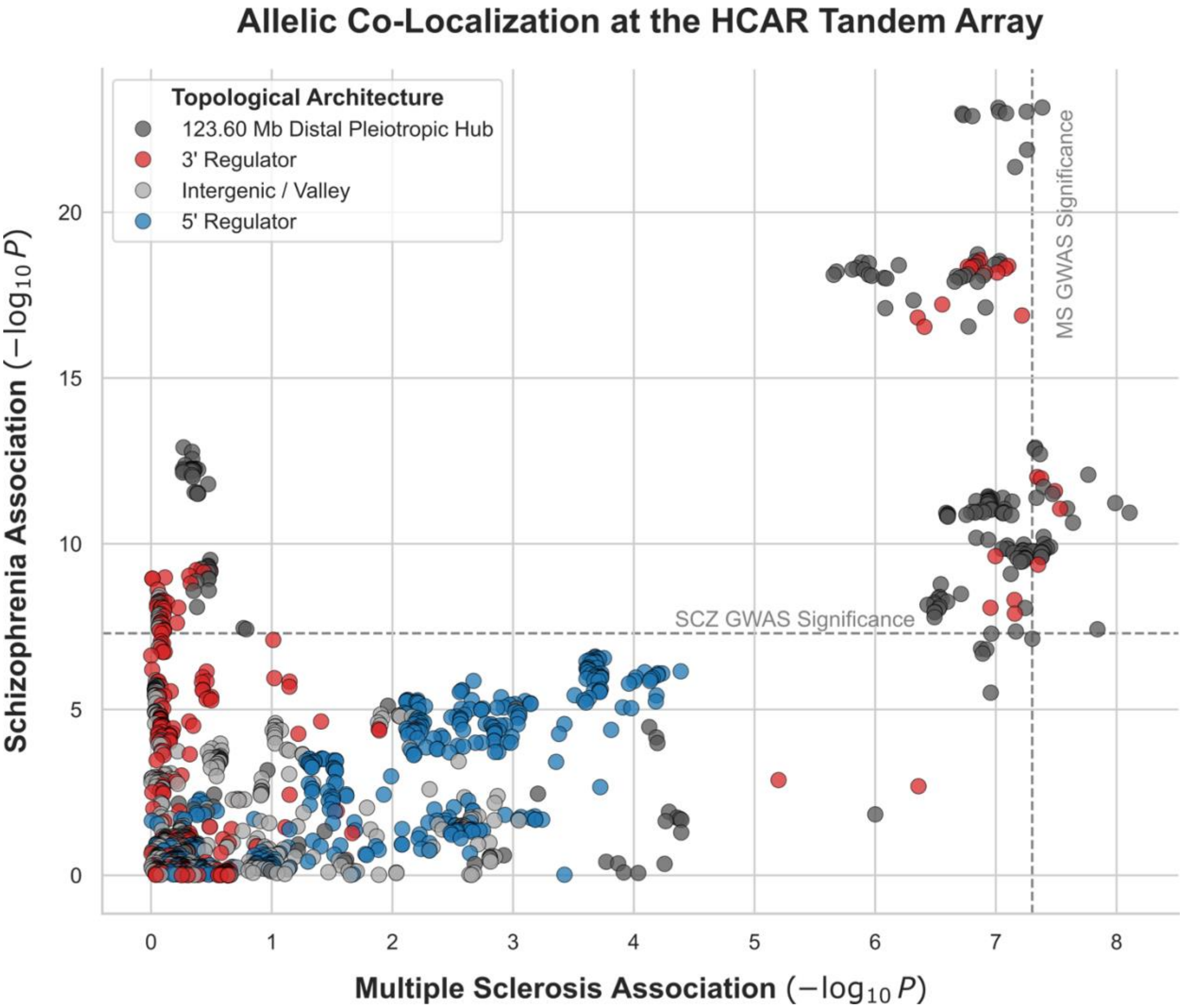


**Supplemental Figure S1. Allelic Co-Localization and Pleiotropic Architecture at the *HCAR* Locus.** A cross-trait significance scatter plot detailing the precise allelic overlap of structural variants spanning the *HCAR* tandem array. The x-axis and y-axis denote the statistical association ($-\log_{10}(P)$) for Multiple Sclerosis (IMSGC Discovery) and Schizophrenia (PGC Wave 3), respectively. Dashed gray lines denote standard genome-wide significance thresholds ($P = 5 \times 10^{-8}$). Variants are color-coded by their physical topological localization: the 5' regulatory enhancer (Blue), the 3' regulatory enhancer (Red), distal hub at 123.60 Mb (Dark Gray), and the intergenic valley spanning *HCAR3* (Light Gray). The plot reveals that while the entire domain exhibits shared variance, true pleiotropic co-localization is overwhelmingly dominated by the 3' regulatory domain and adjacent distal hub at 123.60 Mb. This confirms that the dual-disease susceptibility is driven by the exact same structural fractures potentially acting on the *HCAR* lactate and ketone sensing array.

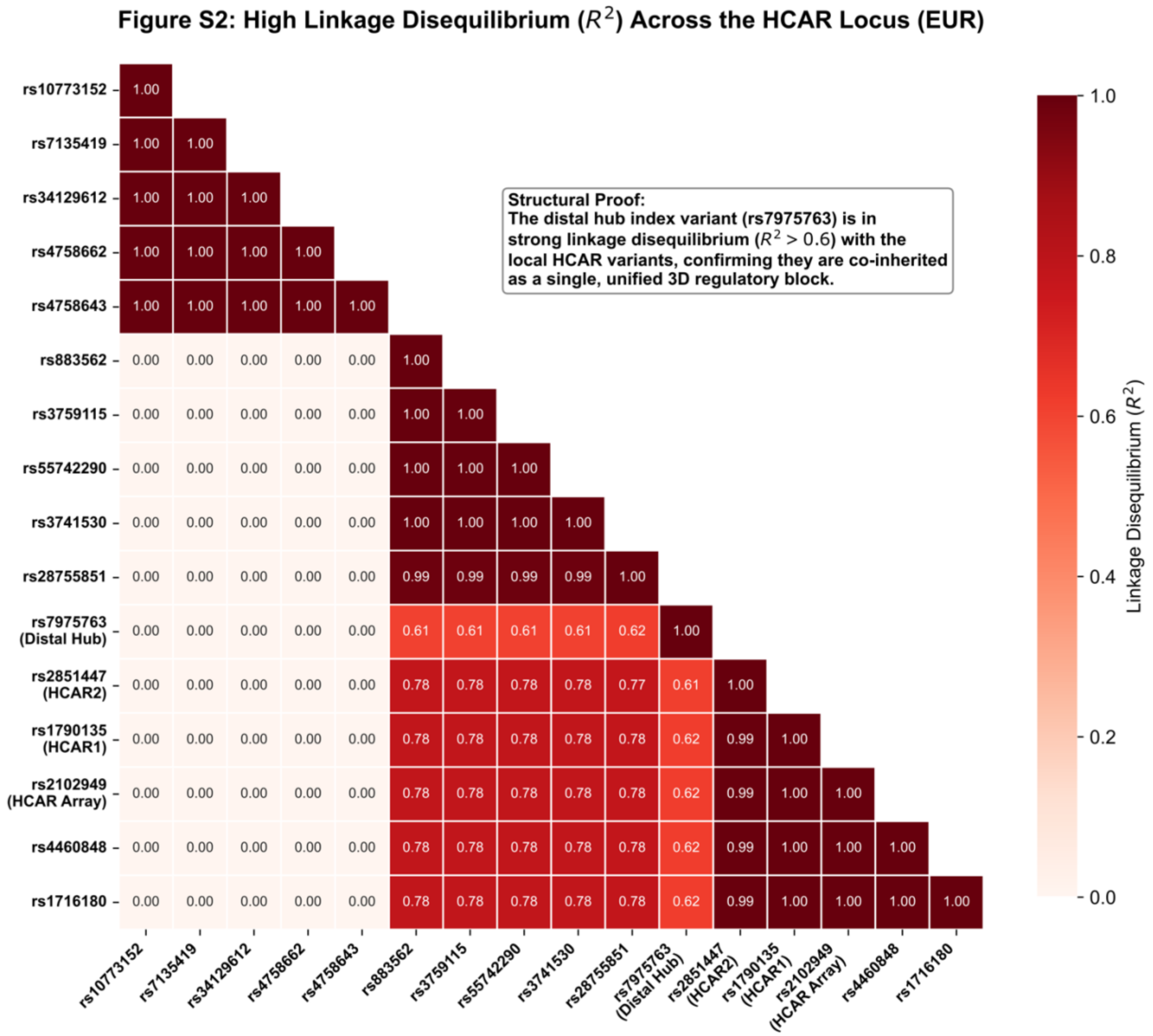


**Supplemental Figure S2. Linkage Disequilibrium (LD) Architecture of the Extended HCAR Locus.** A correlation heatmap displaying the predictive Linkage Disequilibrium ($R^2$) across the 772 kb structural vulnerability block (Chromosome 12: 122.80–123.70 Mb) utilizing the 1000 Genomes European (EUR) reference population. The heatmap explicitly maps the relationship between the 123.60 Mb distal pleiotropic hub (rs7975763, historically misattributed to *PITPNM2*) and the local 5' and 3' regulatory elements flanking the *HCAR* array (e.g., rs1790135, rs2851447). The matrix demonstrates strong linkage disequilibrium ($R^2 > 0.61$, $D' > 0.92$) connecting the distal peak directly to the local HCAR promoter variants. This high correlation mathematically refutes the premise of independent, neighboring association signals (i.e., “LD bleed-over”). Instead, it confirms that the 123.60 Mb distal hub and the local HCAR gene bodies and regulators are co-inherited as a single, unified 3D genomic block, functionally establishing the foundational architecture for the state-dependent multi-gene eQTL regulation observed in activated macrophages.

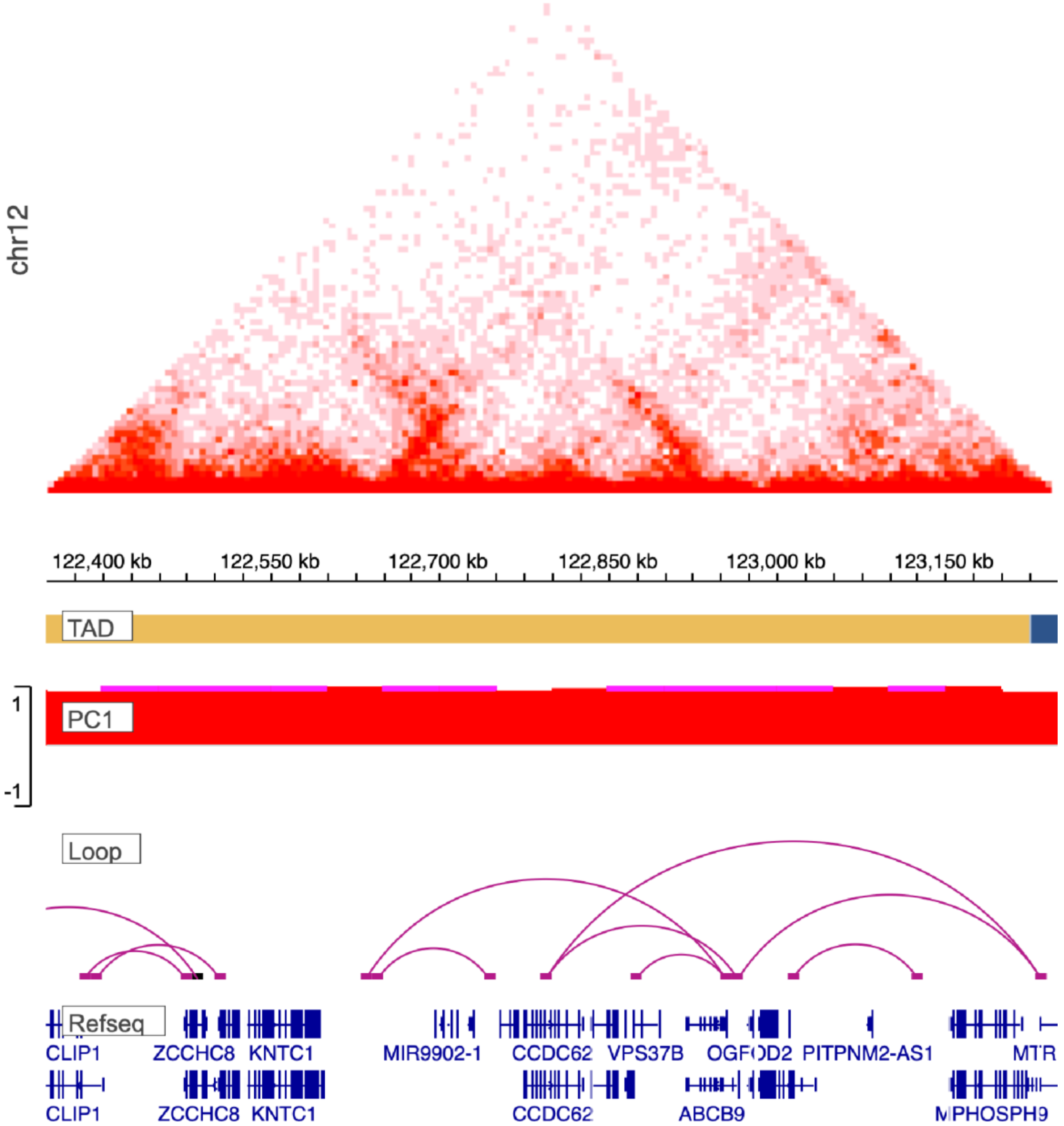


**Supplemental Figure S3. 3D Chromatin Topology of the Extended HCAR Locus in Human Macrophages.** Linear genomic mapping overlaid with 3D chromatin conformation demonstrating the physical interaction landscape of the 122.80–123.70 Mb locus. The primary Schizophrenia and Multiple Sclerosis index variant identified by legacy 1D mapping (rs7975763) resides within the 123.60 Mb distal pleiotropic hub. Crucially, the spatial topology (arcs) confirms that this distal regulatory hub physically bypasses the adjacent *PITPNM2* structural gene body. Instead, it forms direct, long-range enhancer-promoter looping interactions with the local 5' and 3' promoters of the *HCAR1* and *HCAR2* thermodynamic sensors. Spatial architecture was mapped utilizing the 3D Genome Browser (http://3dgenome.fsm.northwestern.edu) rendering *in situ* Hi-C data specifically derived from primary human monocyte-derived macrophages (MDMs) (NCBI GEO Accession: GSE113703). Genomic structural window mapped to hg38 liftover coordinates (chr12:122350000-123250000) to ensure accurate alignment of the distal enhancer and the *HCAR* tandem array.
