## Supplemental Dataset 1 for "State-Dependent 3D Enhancer Architecture Resolves a Shared Schizophrenia and Multiple Sclerosis Ketone and Lactate Sensing Logic Gate": imsgc_ms_vs_scz_cross_reference-widened.py

```
import pandas as pd
import numpy as np

# Define your input files
SCZ_FILE = "Tandem_HCAR_TAD_regulatory_Wave3.tsv"
MS_FILE = "IMSGC_MS_HCAR_Locus_Extracted_widened_window.tsv"

# Define Regulatory Boundaries from the "Mona Lisa" topology & 3D TAD
# Widened to 122.80 to capture the full 5' LD block
REGION_5_PRIME_START = 122_800_000
REGION_5_PRIME_END   = 123_100_000

REGION_3_PRIME_START = 123_300_000
REGION_3_PRIME_END   = 123_500_000

# The Reviewer's Pet Peak (Legacy Assignment to PITPNM2) a.k.a. 123.60 Mb Distal Pleiotropic Hub
REGION_DISTAL_START  = 123_500_001
REGION_DISTAL_END    = 123_700_000

def compare_diseases():
    print("--- Initiating Autoimmune / Psychiatric Cross-Reference (Expanded TAD Edition) ---")
    
    # 1. Load both datasets
    print("Loading Schizophrenia (SCZ) summary stats...")
    df_scz = pd.read_csv(SCZ_FILE, sep='\t')
    
    print("Loading IMSGC Multiple Sclerosis (MS) summary stats...")
    df_ms = pd.read_csv(MS_FILE, sep='\t')
    
    # Clean the IMSGC columns to match our standard pipeline
    # IMSGC Header is: CHR  BP  SNP  A1  A2  N  P  OR
    df_ms = df_ms.rename(columns={
        'BP': 'POS',
        'SNP': 'ID',
        'P': 'PVAL',
        'OR': 'OR_RAW' # Keep this temporarily to calculate BETA
    })
    
    # Convert IMSGC Odds Ratio (OR) to BETA (BETA = ln(OR))
    # This aligns the mathematical direction so we can compare it directly to SCZ Beta
    df_ms['OR_RAW'] = pd.to_numeric(df_ms['OR_RAW'], errors='coerce')
    df_ms['BETA'] = np.log(df_ms['OR_RAW'])
    
    # Ensure POS is numeric for a clean merge
    df_scz['POS'] = pd.to_numeric(df_scz['POS'], errors='coerce')
    df_ms['POS'] = pd.to_numeric(df_ms['POS'], errors='coerce')

    # 2. Merge strictly on the Genomic Coordinate (POS)
    print("Merging datasets on genomic coordinates (POS)...")
    merged_df = pd.merge(
        df_scz, 
        df_ms, 
        on='POS', 
        suffixes=('_SCZ', '_MS'),
        how='inner' # Only keep variants that exist in BOTH datasets
    )
    
    if merged_df.empty:
        print("⚠️ Error: No overlapping coordinates found. Check if genome builds match.")
        return

    # 3. Allele Alignment, Pleiotropy Check, and Architectural Localization
    pleiotropy_results = []
    regulatory_regions = []
    
    for _, row in merged_df.iterrows():
        # SCZ Alleles
        scz_a1 = str(row['A1_SCZ']).upper() # SCZ Effect/Risk Allele
        scz_a2 = str(row['A2_SCZ']).upper()
        scz_beta = float(row['BETA_SCZ'])
        
        # MS Alleles (IMSGC)
        ms_a1 = str(row['A1_MS']).upper()   # MS Effect/Risk Allele
        ms_a2 = str(row['A2_MS']).upper()
        ms_beta = float(row['BETA_MS'])
        
        # Check if the alleles match exactly
        if scz_a1 == ms_a1 and scz_a2 == ms_a2:
            aligned_ms_beta = ms_beta
        # Check if the reference and alternate are flipped between the biobanks
        elif scz_a1 == ms_a2 and scz_a2 == ms_a1:
            aligned_ms_beta = -ms_beta
        else:
            # If alleles don't match at all (e.g., multiallelic sites or strand flips), skip
            aligned_ms_beta = np.nan
            
        # Determine Pleiotropic Direction
        if pd.isna(aligned_ms_beta):
            direction = "Allele Mismatch"
        elif (scz_beta > 0 and aligned_ms_beta > 0) or (scz_beta < 0 and aligned_ms_beta < 0):
            direction = "Shared Risk (Same Direction)"
        else:
            direction = "Antagonistic Pleiotropy (Opposite Direction)"
            
        pleiotropy_results.append(direction)
        
        # Determine Regulatory Architecture Localization
        pos = row['POS']
        if REGION_5_PRIME_START <= pos <= REGION_5_PRIME_END:
            reg_region = "5' Skyscraper (HCAR2)"
        elif REGION_3_PRIME_START <= pos <= REGION_3_PRIME_END:
            reg_region = "3' Skyscraper (HCAR1)"
        elif REGION_DISTAL_START <= pos <= REGION_DISTAL_END:
            reg_region = "123.60 Mb Distal Pleiotropic Hub"
        else:
            reg_region = "Intergenic / Valley"
            
        regulatory_regions.append(reg_region)
        
    merged_df['Pleiotropic_Relationship'] = pleiotropy_results
    merged_df['Regulatory_Region'] = regulatory_regions
    
    # 4. Clean up and sort by SCZ significance to see what MS is doing at the SCZ peaks
    output_cols = [
        'ID_SCZ', 'ID_MS', 'POS', 'Regulatory_Region', 'A1_SCZ', 'A2_SCZ', 
        'PVAL_SCZ', 'BETA_SCZ', 
        'PVAL_MS', 'BETA_MS', 'Pleiotropic_Relationship'
    ]
    
    final_df = merged_df[output_cols].sort_values(by='PVAL_SCZ', ascending=True)
    
    # Export to check your hypothesis
    out_file = "SCZ_vs_IMSGC_MS_Pleiotropy_Analysis_Widened.tsv"
    final_df.to_csv(out_file, sep='\t', index=False)
    
    print(f"\n✅ Cross-reference complete! Found {len(final_df)} overlapping variants.")
    print(f"📁 Exported comparison to: {out_file}")
    
    # Preview the top 5 SCZ hits to see their MS p-values immediately
    print("\n--- TOP 5 SCZ HITS AND THEIR IMSGC MS EFFECTS ---")
    print(final_df[['ID_SCZ', 'ID_MS', 'Regulatory_Region', 'POS', 'PVAL_SCZ', 'PVAL_MS', 'Pleiotropic_Relationship']].head(5).to_string(index=False))

if __name__ == "__main__":
    compare_diseases()
```
