## Supplemental Dataset 1 for "State-Dependent 3D Enhancer Architecture Resolves a Shared Schizophrenia and Multiple Sclerosis Ketone and Lactate Sensing Logic Gate": analyze_scz_vs_ms_severity_py.py

```
import pandas as pd
import numpy as np

# File Paths
SCZ_FILE = "Tandem_HCAR2_HCAR1_wide_regulatory.tsv"
MS_SEV_FILE = "IMSGC_MS_SEVERE_HCAR_Locus_Extracted.tsv"

# Genomic Topology
REGION_5_PRIME_START = 122_900_000
REGION_5_PRIME_END   = 123_100_000
REGION_3_PRIME_START = 123_300_000
REGION_3_PRIME_END   = 123_500_000

def analyze_severity_overlap():
    print("--- Initiating SCZ vs. MS Severity Cross-Reference ---")
    
    # 1. Load Data
    df_scz = pd.read_csv(SCZ_FILE, sep='\t')
    df_ms = pd.read_csv(MS_SEV_FILE, sep='\t')
    
    # Ensure POS is numeric for a clean merge
    df_scz['POS'] = pd.to_numeric(df_scz['POS'], errors='coerce')
    df_ms['POS'] = pd.to_numeric(df_ms['POS'], errors='coerce')
    
    # 2. Merge on POS
    # Using 'SNP' or 'ID' column if needed, but POS is primary
    merged_df = pd.merge(df_scz, df_ms, on='POS', suffixes=('_SCZ', '_MS'))
    
    # 3. Annotate Regulatory Regions
    regions = []
    for pos in merged_df['POS']:
        if REGION_5_PRIME_START <= pos <= REGION_5_PRIME_END:
            regions.append("5' Skyscraper (HCAR2)")
        elif REGION_3_PRIME_START <= pos <= REGION_3_PRIME_END:
            regions.append("3' Skyscraper (HCAR1)")
        else:
            regions.append("Intergenic / Valley")
    merged_df['Regulatory_Region'] = regions
    
    # 4. Annotate Direction of Pleiotropy
    # Because both files have a 'BETA' column, pandas suffixed them to 'BETA_SCZ' and 'BETA_MS'
    merged_df['Pleiotropic_Relationship'] = np.where(
        (merged_df['BETA_SCZ'] * merged_df['BETA_MS']) > 0,
        "Shared Risk (Same Direction)",
        "Antagonistic Pleiotropy (Opposite Direction)"
    )
    
    # 5. Corrected Output Columns based on exact header names from the SCZ and MS files
    # SCZ uses 'ID' and 'PVAL'. MS uses 'SNP' and 'P'.
    output_cols = [
        'ID', 'SNP', 'POS', 'Regulatory_Region', 
        'PVAL', 'BETA_SCZ', 'P', 'BETA_MS', 'Pleiotropic_Relationship'
    ]
    
    # Filter columns that actually exist to avoid KeyError
    final_cols = [c for c in output_cols if c in merged_df.columns]
    
    # Sort by the actual SCZ P-value column name ('PVAL')
    final_df = merged_df[final_cols].sort_values(by='PVAL')
    final_df.to_csv("SCZ_vs_MS_Severity_Analysis.tsv", sep='\t', index=False)
    
    print(f"✅ Analysis complete! Found {len(final_df)} overlapping variants.")
    print("Results saved to: SCZ_vs_MS_Severity_Analysis.tsv")

if __name__ == "__main__":
    analyze_severity_overlap()
```
