## Supplemental Dataset 1 for "State-Dependent 3D Enhancer Architecture Resolves a Shared Schizophrenia and Multiple Sclerosis Ketone and Lactate Sensing Logic Gate": imsgc_severity_extractor_bash.sh

```
#!/bin/bash

# Define Target Locus (hg19 / GRCh37)
CHROM="12"
START_BP=122850000
END_BP=123515000

# Explicitly targeting the un-tarred MS Severity TSV
INPUT_FILE="imsgc_mssev_discovery.tsv" 
OUTPUT_FILE="IMSGC_MS_SEVERE_HCAR_Locus_Extracted.tsv"

echo "--- Initiating IMSGC Severity Targeted Extraction ---"
echo "Target File: $INPUT_FILE"
echo "Locus: chr$CHROM : $START_BP - $END_BP"

# Use 'cat' since the file is un-tarred and uncompressed plain text (.tsv)
cat "$INPUT_FILE" | awk -v chr="$CHROM" -v start="$START_BP" -v end="$END_BP" '
BEGIN {
    OFS="\t"; 
    # Fallback columns (readme.txt specifies POS is in column 3)
    chr_col = 1; pos_col = 3;
}
NR==1 {
    # Dynamically find the Chromosome and Position columns in the IMSGC header
    for (i=1; i<=NF; i++) {
        if ($i == "CHR" || $i == "CHROM" || $i == "chr") chr_col = i;
        if ($i == "BP" || $i == "POS" || $i == "pos") pos_col = i;
    }
    $1=$1 
    print $0 
    next
}
($chr_col == chr || $chr_col == "chr"chr) && $pos_col >= start && $pos_col <= end {
    $1=$1 
    print $0
}
' > "$OUTPUT_FILE"

HIT_COUNT=$(wc -l < "$OUTPUT_FILE" | tr -d ' ')
HIT_COUNT=$((HIT_COUNT - 1))

echo "✅ Extraction Complete!"
echo "Isolated $HIT_COUNT severity variants within the HCAR topological domain."
echo "Data saved to: $OUTPUT_FILE"
```
