## Supplemental Dataset 1 for "State-Dependent 3D Enhancer Architecture Resolves a Shared Schizophrenia and Multiple Sclerosis Ketone and Lactate Sensing Logic Gate": readme.txt

# Multiple Sclerosis Severity GWAS Summary Statistics

Authors: International Multiple Sclerosis Genetics Consortium, MultipleMS

Date last updated: April 12, 2023
Written by Adil Harroud

## Data Usage Agreement

By registering to use these data, you have agreed to the following terms:

- You will not redistribute, sell or otherwise distribute these data to third parties.
- You will include the IMSGC as a co-author or acknowledge the IMSGC in any publication that proceeds from the use of these data.
- You will send the International Multiple Sclerosis Genetics Consortium (IMSGC) a copy of any paper published by you using these data (in part or in full).
- These data and results may not be used in attempts to identify individual study participants.
- The contact details you have provided may be used to contact you in the context of this data request and for any useful follow-up.

## Analysis Details

The analysis details are described in the Nature publication entitled “Novel genetic locus and CNS resilience modulate multiple sclerosis severity” and can be found at the following DOI: https://doi.org/10.1038/s41586-023-06250-x
## File Contents

The file contains the following columns:

- CHR	: chromosome number
- SNP	: variant rs number
- POS	: Base pair position (GRCh37)
- A1	: Alternate (effect) allele
- A2	: Reference (non-effect) allele
- AF1	: Alternate allele frequency
- BETA	: linear regression coefficient, that is per-allele effect size on the age-related multiple sclerosis severity score following rank-based inverse normal transformation 
- SE	: Standard error of the regression coefficient
- P		: P-value of the null hypothesis that the coefficient is equal to zero
- Rsq	: Minimac imputation quality score
