## Supplemental Dataset 1 for "State-Dependent 3D Enhancer Architecture Resolves a Shared Schizophrenia and Multiple Sclerosis Ketone and Lactate Sensing Logic Gate": README.docx

**Supplemental Dataset 1: IMSGC Multiple Sclerosis Susceptibility and Severity Mining**

**Study:** State-Dependent 3D Enhancer Architecture Resolves a Shared Schizophrenia and Multiple Sclerosis Ketone and Lactate Sensing Logic Gate

**Author:** Bryan A. Krantz

**Overview**

This repository contains the extraction pipelines, manual curation scripts, and the targeted genomic output files used to map the *HCAR* tandem array against Multiple Sclerosis (MS).

To test the hypothesis that the *HCAR1/HCAR2* mutational skyscrapers act strictly as a neuro-immune "ignition switch," this dataset evaluates two distinct MS phenotypes:

1. **MS Susceptibility (Discovery):** Does the SCZ structural variance dictate whether a patient develops MS?
2. **MS Severity:** Does the same structural variance dictate the downstream velocity of demyelination and neural decay?

**⚠️ Important Note on Data Access**

The master, full-genome summary statistic files utilized in this pipeline are restricted to protect patient privacy and are governed by the International Multiple Sclerosis Genetics Consortium (IMSGC).

- discovery_metav3.0.meta.gz (Susceptibility)
- imsgc_mssev_discovery.tar.gz / .tsv (Severity)

**These massive master files have been deleted from this public repository to comply with data-sharing agreements.** Researchers wishing to replicate the extraction from scratch must request access to the primary datasets directly via the IMSGC website (http://imsgc.net).

This repository provides the custom extraction scripts and the resulting lightweight, locus-specific files (restricted exactly to Chromosome 12: 122.80 Mb – 123.70 Mb; hg19) utilized for downstream cross-trait analyses.

**The Psychiatric Baseline Data (Tandem_HCAR_TAD_regulatory_Wave3.tsv)**

Throughout this directory, the MS extractions are cross-referenced against the baseline Schizophrenia architecture. This SCZ baseline file (Tandem_HCAR_TAD_regulatory_Wave3.tsv) was extracted from the massive PGC Wave 3 Schizophrenia meta-analysis using the following wide regulatory sweep (Chromosome 12: 122.80 Mb – 123.70 Mb; hg19) using bash-HCAR2-HCAR1-widened-regulatory-extractor.sh:

#!/bin/bash

\# Wide Regulatory Sweep of HCAR1/2/3 and PITPNM2 TAD Locus (122.80 Mb - 123.70 Mb)

zgrep -v "^##" PGC3_SCZ_wave3.primary.autosome.public.v3.vcf.tsv.gz | awk 'NR==1 {print} $1=="12" && $3>=122800000 && $3<=123700000 {print}' > Tandem_HCAR_TAD_regulatory_Wave3.tsv

For full details on the primary SCZ baseline extraction refer to the primary Zenodo deposition (DOI: 10.5281/zenodo.20930404).

**Directory Structure and File Manifest**

**1. discovery_metav3.0.meta/ (MS Susceptibility)**

This directory contains the pipeline determining if the structural variance drives disease onset.

- **Extraction Scripts:**
  - imsgc_ms_extractor_bash_delimiter_fix_widened_window.sh: The standard Unix pipeline utilizing awk and zgrep to slice the master .gz file down to the *HCAR* topological boundaries. The script was modified accounting for dynamic header formatting and irregular tab/space delimiters in the raw IMSGC file.
- **Reference Input:**
  - Tandem_HCAR_TAD_regulatory_Wave3.tsv: The baseline PGC3 Schizophrenia reference data imported for pleiotropy mapping (detailed above).
- **Analysis and Outputs:**
  - IMSGC_MS_HCAR_Locus_Extracted_widened_window.tsv: The lightweight, locus-specific MS GWAS extraction.
  - imsgc_ms_vs_scz_cross_reference-widened.py: The Python script that merges the MS locus with the SCZ baseline, algorithmically aligning the risk alleles and calculating the pleiotropic direction (Shared Risk vs. Antagonistic).
  - *(Note: The output from this script provides the primary data for the Figure 1 dual-ignition locus zoom).*

**2. imsgc_mssev_discovery/ (MS Severity Negative Control)**

This directory contains the pipeline utilized as a negative control to prove the *HCAR* locus does not govern downstream demyelination velocity.

- **Extraction Scripts:**
  - imsgc_severity_extractor_widened_window_bash.sh: Slices the uncompressed MS severity dataset (imsgc_mssev_discovery.tsv) to the *HCAR* boundaries.
- **Reference Input:**
  - readme.txt: Original data dictionary and column key provided by the IMSGC.
  - Tandem_HCAR_TAD_regulatory_Wave3.tsv: The baseline PGC3 Schizophrenia reference data.
- **Analysis and Outputs:**
  - IMSGC_MS_SEVERE_HCAR_Locus_Extracted_widened_window.tsv: The localized severity summary statistics.
  - analyze_scz_vs_ms_severity_widened_window.py: The Python script that merges the severity data against the SCZ baseline to check for shared variance.
  - SCZ_vs_MS_Severity_Analysis_widened_window.tsv: The final output matrix demonstrating total statistical noise (lack of association), confirming the locus acts strictly as an ignition switch. This file serves as a direct input for the Figure 2 Neuro-Immune Heatmap.
