## Supplemental Dataset 2 for "State-Dependent 3D Enhancer Architecture Resolves a Shared Schizophrenia and Multiple Sclerosis Ketone and Lactate Sensing Logic Gate": extract_ukbb_hcar_locus.sh

```
#!/bin/bash

if [ -z "$1" ]; then
  echo "⚠️ Usage: ./extract_ukbb_hcar_locus.sh <ukbb_summary_stats.tsv.bgz>"
  exit 1
fi

INPUT_FILE=$1
OUTPUT_FILE="UKBB_HCAR_Locus_Extracted_${INPUT_FILE%.tsv*}_chr12.tsv"

CHROM="12"
START_BP=122850000
END_BP=123515000

echo "--- Initiating Targeted Extraction ---"
echo "Target File: $INPUT_FILE"
echo "Locus: chr$CHROM : $START_BP - $END_BP"

# 1. Grab the header
gzip -dc "$INPUT_FILE" | head -n 1 > "$OUTPUT_FILE"

echo "Scanning for structural variance within the HCAR regulatory domains..."

# 2. Extract the locus
echo "Scanning for structural variance within the HCAR regulatory domains..."
gzip -dc "$INPUT_FILE" | awk -v chr="$CHROM" -v start="$START_BP" -v end="$END_BP" 'BEGIN {FS="\t"; OFS="\t"} ($1 == chr || $1 == "chr"chr) && $2 >= start && $2 <= end {print $0}' >> "$OUTPUT_FILE"

# 3. Calculate hits (stripping Mac's invisible spaces)
HIT_COUNT=$(wc -l < "$OUTPUT_FILE" | tr -d ' ')
HIT_COUNT=$((HIT_COUNT - 1))

echo "✅ Extraction Complete!"
echo "Isolated $HIT_COUNT variants within the HCAR topological domain."
echo "Data saved to: $OUTPUT_FILE"
```
