## Supplemental Dataset 2 for "State-Dependent 3D Enhancer Architecture Resolves a Shared Schizophrenia and Multiple Sclerosis Ketone and Lactate Sensing Logic Gate": pan_ukbb_manifest_autoimmune_filter.py

```
import pandas as pd
import os

def mine_pan_ukbb_manifest():
    print("--- Hunting for Autoimmune GWAS Targets in Pan-UKBB ---")
    
    manifest_file = "Pan-UK Biobank phenotype manifest - phenotype_manifest.csv"
    
    if not os.path.exists(manifest_file):
        print(f"⚠️ Error: Could not find '{manifest_file}'.")
        return

    # 1. Load the manifest
    try:
        df = pd.read_csv(manifest_file, low_memory=False)
        print(f"Loaded massive manifest with {len(df)} phenotypes.")
    except Exception as e:
        print(f"⚠️ Error reading manifest: {e}")
        return

    # Ensure column names are lowercase string for easier targeting
    df.columns = [str(col).lower().strip() for col in df.columns]
    
    # 2. Define our Autoimmune Targets (ICD-10 Codes & Keywords)
    # The Pan-UKBB often uses the 'phenocode' column for ICD-10, and 'description' or 'trait_name' for text.
    target_icd10 = ['K50', 'M05', 'M06', 'M32', 'G35', 'L40']
    target_keywords = ['crohn', 'rheumatoid', 'lupus', 'multiple sclerosis', 'psoriasis']
    
    # Identify the correct description column (varies slightly by manifest version)
    desc_col = 'description' if 'description' in df.columns else 'trait_name' if 'trait_name' in df.columns else None
    pheno_col = 'phenocode' if 'phenocode' in df.columns else None
    
    if not desc_col or not pheno_col:
        print("⚠️ Warning: Could not identify standard 'description' or 'phenocode' columns. Check the manifest headers.")
        print(f"Available columns: {list(df.columns)}")
        return

    # 3. Apply the Filters
    # Fill NAs to prevent regex/string matching errors
    df[desc_col] = df[desc_col].fillna('')
    df[pheno_col] = df[pheno_col].fillna('')

    # Filter by Keyword (case-insensitive)
    keyword_mask = df[desc_col].str.contains('|'.join(target_keywords), case=False)
    
    # Filter by ICD-10 Code
    icd_mask = df[pheno_col].str.contains('|'.join(target_icd10), case=False)
    
    # Combine masks
    target_df = df[keyword_mask | icd_mask].copy()

    # 4. Filter for Power (Optional but highly recommended)
    # You want phenotypes with a decent number of cases, otherwise the GWAS is just noise.
    if 'n_cases_white_british' in df.columns:
        # Assuming we want to match the predominant ancestry of PGC3 for apples-to-apples mapping
        target_df = target_df[target_df['n_cases_white_british'] > 500]
    elif 'n_cases_eur' in df.columns:
        target_df = target_df[target_df['n_cases_eur'] > 500]

    # 5. Format the Output
    # We want the disease name, the code, the number of cases, and the exact file/wget link
    output_cols = [pheno_col, desc_col]
    
    # Add case counts and download links if they exist in the columns
    for col in ['n_cases_eur', 'n_cases_white_british', 'wget_command', 'aws_link', 'file_name', 'tabix_link']:
        if col in df.columns:
            output_cols.append(col)
            
    final_hits = target_df[output_cols]
    
    # Export the hit-list
    output_csv = "Autoimmune_Target_GWAS_Files.csv"
    final_hits.to_csv(output_csv, index=False)
    
    print(f"\n✅ Success! Narrowed down to {len(final_hits)} high-powered autoimmune targets.")
    print(f"📁 Target manifest saved to: {output_csv}")
    print("\n--- PREVIEW OF TOP HITS ---")
    print(final_hits[[pheno_col, desc_col]].head(10).to_string(index=False))

if __name__ == "__main__":
    mine_pan_ukbb_manifest()
```
