## Supplemental Dataset 2 for "State-Dependent 3D Enhancer Architecture Resolves a Shared Schizophrenia and Multiple Sclerosis Ketone and Lactate Sensing Logic Gate": psoriasis_vs_scz_cross_reference.py

```
import pandas as pd
import numpy as np

# Define your input files
SCZ_FILE = "Tandem_HCAR2_HCAR1_wide_regulatory.tsv"
PSORIASIS_FILE = "UKBB_HCAR_Locus_Extracted_icd10-L40-both_sexes_chr12.tsv"

# Define Regulatory Boundaries from the "Mona Lisa" topology
REGION_5_PRIME_START = 122_900_000
REGION_5_PRIME_END   = 123_100_000

REGION_3_PRIME_START = 123_300_000
REGION_3_PRIME_END   = 123_500_000

def compare_diseases():
    print("--- Initiating Autoimmune / Psychiatric Cross-Reference (Psoriasis) ---")
    
    # 1. Load both datasets
    print("Loading Schizophrenia (SCZ) summary stats...")
    df_scz = pd.read_csv(SCZ_FILE, sep='\t')
    
    print("Loading Psoriasis summary stats...")
    df_psoriasis = pd.read_csv(PSORIASIS_FILE, sep='\t')
    
    # Ensure POS is numeric for a clean merge
    df_scz['POS'] = pd.to_numeric(df_scz['POS'], errors='coerce')
    df_psoriasis['POS'] = pd.to_numeric(df_psoriasis['POS'], errors='coerce')

    # 2. Merge strictly on the Genomic Coordinate (POS)
    print("Merging datasets on genomic coordinates (POS)...")
    merged_df = pd.merge(
        df_scz, 
        df_psoriasis, 
        on='POS', 
        suffixes=('_SCZ', '_PSORIASIS'),
        how='inner' # Only keep variants that exist in BOTH datasets
    )
    
    if merged_df.empty:
        print("⚠️ Error: No overlapping coordinates found. Check if genome builds match.")
        return

    # 3. Allele Alignment, Pleiotropy Check, and Architectural Localization
    pleiotropy_results = []
    regulatory_regions = []
    
    for _, row in merged_df.iterrows():
        # SCZ Alleles
        scz_a1 = str(row['A1_SCZ']).upper() # SCZ Effect/Risk Allele
        scz_a2 = str(row['A2_SCZ']).upper()
        scz_beta = float(row['BETA_SCZ'])
        
        # Psoriasis Alleles
        psoriasis_a1 = str(row['A1_PSORIASIS']).upper()   # Psoriasis Effect/Risk Allele
        psoriasis_a2 = str(row['A2_PSORIASIS']).upper()
        psoriasis_beta = float(row['BETA_PSORIASIS'])
        
        # Check if the alleles match exactly
        if scz_a1 == psoriasis_a1 and scz_a2 == psoriasis_a2:
            aligned_psoriasis_beta = psoriasis_beta
        # Check if the reference and alternate are flipped between the biobanks
        elif scz_a1 == psoriasis_a2 and scz_a2 == psoriasis_a1:
            aligned_psoriasis_beta = -psoriasis_beta
        else:
            # If alleles don't match at all (e.g., multiallelic sites or strand flips), skip
            aligned_psoriasis_beta = np.nan
            
        # Determine Pleiotropic Direction
        if pd.isna(aligned_psoriasis_beta):
            direction = "Allele Mismatch"
        elif (scz_beta > 0 and aligned_psoriasis_beta > 0) or (scz_beta < 0 and aligned_psoriasis_beta < 0):
            direction = "Shared Risk (Same Direction)"
        else:
            direction = "Antagonistic Pleiotropy (Opposite Direction)"
            
        pleiotropy_results.append(direction)
        
        # Determine Regulatory Architecture Localization
        pos = row['POS']
        if REGION_5_PRIME_START <= pos <= REGION_5_PRIME_END:
            reg_region = "5' Skyscraper (HCAR2)"
        elif REGION_3_PRIME_START <= pos <= REGION_3_PRIME_END:
            reg_region = "3' Skyscraper (HCAR1)"
        else:
            reg_region = "Intergenic / Valley"
            
        regulatory_regions.append(reg_region)
        
    merged_df['Pleiotropic_Relationship'] = pleiotropy_results
    merged_df['Regulatory_Region'] = regulatory_regions
    
    # 4. Clean up and sort by SCZ significance to see what Psoriasis is doing at the SCZ peaks
    output_cols = [
        'ID_SCZ', 'POS', 'Regulatory_Region', 'A1_SCZ', 'A2_SCZ', 
        'PVAL_SCZ', 'BETA_SCZ', 
        'PVAL_PSORIASIS', 'BETA_PSORIASIS', 'Pleiotropic_Relationship'
    ]
    
    final_df = merged_df[output_cols].sort_values(by='PVAL_SCZ', ascending=True)
    
    # Export to check your hypothesis
    out_file = "SCZ_vs_Psoriasis_Pleiotropy_Analysis.tsv"
    final_df.to_csv(out_file, sep='\t', index=False)
    
    print(f"\n✅ Cross-reference complete! Found {len(final_df)} overlapping variants.")
    print(f"📁 Exported comparison to: {out_file}")
    
    # Preview the top 5 SCZ hits to see their Psoriasis p-values immediately
    print("\n--- TOP 5 SCZ HITS AND THEIR PSORIASIS EFFECTS ---")
    print(final_df[['ID_SCZ', 'Regulatory_Region', 'POS', 'PVAL_SCZ', 'PVAL_PSORIASIS', 'Pleiotropic_Relationship']].head(5).to_string(index=False))

if __name__ == "__main__":
    compare_diseases()
```
