## Supplemental Dataset 2 for "State-Dependent 3D Enhancer Architecture Resolves a Shared Schizophrenia and Multiple Sclerosis Ketone and Lactate Sensing Logic Gate": README.docx

**Supplemental Dataset 2: Pan-UK Biobank Autoimmune Mining and Cross-Trait Pleiotropy Analysis**

**Study:** State-Dependent 3D Enhancer Architecture Resolves a Shared Schizophrenia and Multiple Sclerosis Ketone and Lactate Sensing Logic Gate

**Author:** Bryan A. Krantz

**Overview**

This repository contains the data mining pipeline, raw extraction logs, and cross-reference analysis scripts utilized to establish the "Neuro-Immune Bifurcation" of the *HCAR* locus.

To prove that the *HCAR1/HCAR2* tandem structural fracture is a highly specific driver of neuro-immune pathology (Schizophrenia and Multiple Sclerosis) rather than a generic systemic inflammation hotspot, the SCZ index variants were cross-referenced against classic autoimmune diseases. High-powered GWAS summary statistics were systematically mined from the Pan-UK Biobank (Pan-UKBB) multi-ancestry meta-analysis.

The resulting outputs from this pipeline (SCZ_vs_[Disease]_Pleiotropy_Analysis _Widened_Window.tsv) serve as the direct empirical inputs for the Negative Control Heatmap (**Figure 2**).

**Directory Structure and Workflow**

**1. Parent Directory: The Manifest Filter**

The root of this dataset handles the programmatic triage of the Pan-UKBB.

- Pan-UK Biobank phenotype manifest - phenotype_manifest.csv: The raw, massive metadata file detailing all available phenotypes in the biobank.
- pan_ukbb_manifest_autoimmune_filter.py: A custom Python script that parses the manifest using targeted ICD-10 codes (e.g., K50, L40) and Phecodes (e.g., 714, 695.4) to isolate high-powered autoimmune GWAS datasets.
- Autoimmune_Target_GWAS_Files.csv: The resulting target list containing the direct wget URLs used to download the multi-gigabyte .tsv.bgz summary statistics files into their respective subdirectories.

**2. Disease-Specific Subdirectories**

The analysis is modularized into distinct subdirectories for each evaluated phenotype:

- crohns_K50/ (Crohn's Disease)
- lupus_phecode_695.4/ (Systemic Lupus Erythematosus)
- psoriasis_L40/ (Psoriasis)
- rheumatoid_arthritis_and_other_inflamatory_polyarthropathies_phecode_714/ (Rheumatoid Arthritis)
- multiple_sclerosis_phecode_355/ (UKBB MS Cohort Validation)

**3. Pipeline Execution (Within Each Subdirectory)**

Inside each phenotype folder, a standardized three-step pipeline was executed:

**Step A: Raw Data Acquisition**

- [phenotype_code]-both_sexes.tsv.bgz: The raw, whole-genome summary statistics downloaded directly from Pan-UKBB. *(Note: Due to file size constraints, these raw whole-genome files may be excluded from the final repository upload but can be regenerated using the wget links in the parent directory).*

**Step B: Locus Extraction and Harmonization**

- extract_ukbb_hcar_locus_widened_window_ver3.sh: A highly optimized Unix Bash script that utilizes zgrep and awk to slice the massive .bgz file down to the exact Chromosome 12 topological boundaries of the *HCAR* locus (Chromosome 12: 122.80 Mb – 123.70 Mb; hg19). Crucially, this script also dynamically renames and formats the UKBB columns to perfectly match the PGC Wave 3 standard (e.g., constructing proxy rsIDs and converting -log_10_ *P* to raw *P*-values).
- UKBB_HCAR_Locus_Extracted_[code]_chr12 _widened_window.tsv: The resulting lightweight, extracted target locus.

**Step C: Psychiatric Cross-Referencing**

- Tandem_HCAR_TAD_regulatory_Wave3.tsv: The baseline PGC3 Schizophrenia reference data (copied into each folder for local execution).
- Python widened window .py script: The core analytical Python script. It merges the extracted UKBB data with the SCZ data based on exact genomic coordinates. It algorithmically aligns the alleles (accounting for strand flips or reference/alternate inversions between the biobanks) and calculates the pleiotropic directionality by multiplying the effect sizes (Betas).
- SCZ_vs_[Disease]_Pleiotropy_Analysis_Widened_Window.tsv: The final output matrix. It annotates each variant with its topological regulatory domain (5' regulatory region, 3' regulatory region, distal hub at 123.60 Mb, or Intergenic) and provides the aligned P-values and effect directions for immediate ingestion by the Figure 2 heatmap generator.
