## Supplemental Dataset 2 for "State-Dependent 3D Enhancer Architecture Resolves a Shared Schizophrenia and Multiple Sclerosis Ketone and Lactate Sensing Logic Gate": extract_ukbb_hcar_locus_ver2.sh

```
#!/bin/bash

if [ -z "$1" ]; then
  echo "⚠️ Usage: ./extract_ukbb_hcar_locus.sh <ukbb_summary_stats.tsv.bgz>"
  exit 1
fi

INPUT_FILE=$1
OUTPUT_FILE="UKBB_HCAR_Locus_Extracted_${INPUT_FILE%.tsv*}_chr12.tsv"

CHROM="12"
START_BP=122850000
END_BP=123515000

echo "--- Initiating Targeted Extraction ---"
echo "Target File: $INPUT_FILE"
echo "Locus: chr$CHROM : $START_BP - $END_BP"

# 1. Grab and Format the Header (PGC Wave 3 Standard)
printf "CHROM\tID\tPOS\tA1\tA2\tPVAL\tBETA\tSE\tNEG_LOG10_P\tLOW_CONFIDENCE\n" > "$OUTPUT_FILE"

# 2. Extract the locus and translate UKBB format to PGC format
echo "Scanning for structural variance within the HCAR regulatory domains..."
gzip -dc "$INPUT_FILE" | awk -v chr="$CHROM" -v start="$START_BP" -v end="$END_BP" '
BEGIN {FS="\t"; OFS="\t"}
NR > 1 && ($1 == chr || $1 == "chr"chr) && $2 >= start && $2 <= end {
    # Convert -log10(P) back to standard P-value (UKBB Column 9)
    pval = 10^(-$9)
    
    # Create a proxy rsID since UKBB drops them (Format: chr:pos_ref_alt)
    proxy_id = "chr" $1 ":" $2 "_" $3 "_" $4
    
    # Print in standard PGC format: CHROM, ID, POS, A1(alt), A2(ref), PVAL, plus the original stats
    print $1, proxy_id, $2, $4, $3, pval, $7, $8, $9, $10
}' >> "$OUTPUT_FILE"

# 3. Calculate hits (stripping Mac's invisible spaces)
HIT_COUNT=$(wc -l < "$OUTPUT_FILE" | tr -d ' ')
HIT_COUNT=$((HIT_COUNT - 1))

echo "✅ Extraction Complete!"
echo "Isolated $HIT_COUNT variants within the HCAR topological domain."
echo "Data saved to: $OUTPUT_FILE"
```
