## Supplemental Dataset 3 for "State-Dependent 3D Enhancer Architecture Resolves a Shared Schizophrenia and Multiple Sclerosis Ketone and Lactate Sensing Logic Gate": architectural_enhancer_audit.py

```
import pandas as pd

INPUT_FILE = "Macrophage_SCZ_MS_Smoking_Gun.tsv"

def audit_architecture():
    print("--- Initiating 3D Architectural Enhancer Audit ---\n")
    
    try:
        df = pd.read_csv(INPUT_FILE, sep='\t')
    except FileNotFoundError:
        print(f"❌ Error: Could not find {INPUT_FILE}")
        return
        
    # Let's fix the nomenclature right now so we stop validating the reviewer
    df['Regulatory_Region'] = df['Regulatory_Region'].replace(
        "Distal Enhancer (PITPNM2 / Reviewer's Peak)", 
        "123.60 Mb Distal Pleiotropic Hub"
    )
    
    print("Separating Local Skyscrapers from the Distal Hub...\n")
    
    # Group by Region, Gene, and Consequence to see the spatial breakdown
    summary = df.groupby(['Regulatory_Region', 'Target_Gene', 'Biological_Consequence']).size().reset_index(name='Total_SNPs')
    
    # Sort for readability
    summary = summary.sort_values(by=['Target_Gene', 'Regulatory_Region'])
    
    print("=====================================================================")
    print("🧬 SPATIAL ARCHITECTURE OF MACROPHAGE TRANSCRIPTOMIC CONTROL")
    print("=====================================================================")
    print(summary.to_string(index=False))
    print("=====================================================================\n")

    print("--- Reviewer's Pet SNP (rs7975763) Spatial Target Check ---")
    pet_snp_data = df[df['rsID'] == 'rs7975763']
    if not pet_snp_data.empty:
        for _, row in pet_snp_data.iterrows():
            print(f"rs7975763 physically acts from: {row['Regulatory_Region']}")
            print(f"   -> Regulates: {row['Target_Gene']} ({row['Biological_Consequence']})")
    else:
        print("rs7975763 not found in the significant eQTL pool.")

if __name__ == "__main__":
    audit_architecture()
```
