## Supplemental Dataset 3 for "State-Dependent 3D Enhancer Architecture Resolves a Shared Schizophrenia and Multiple Sclerosis Ketone and Lactate Sensing Logic Gate": audit_directionality.py

```
import pandas as pd

EBI_FILE = "EBI_Immune_eQTL_Targeted_DeepSweep_Hits.tsv"
PLEIO_FILE = "SCZ_vs_IMSGC_MS_Pleiotropy_Analysis_Widened.tsv"

# The specific SNPs we want to audit
TARGET_SNPS = ["rs7975763", "rs1790135"]

def audit_directionality():
    print("--- Initiating Beta / Allele Directionality Audit ---\n")
    
    # 1. Load GWAS Data for the targets
    df_pleio = pd.read_csv(PLEIO_FILE, sep='\t')
    gwas_targets = df_pleio[df_pleio['ID_SCZ'].isin(TARGET_SNPS)]
    
    gwas_dict = {}
    for _, row in gwas_targets.iterrows():
        gwas_dict[row['ID_SCZ']] = {
            'A1_Risk': str(row['A1_SCZ']).upper(),
            'A2_Ref': str(row['A2_SCZ']).upper(),
            'SCZ_Beta': float(row['BETA_SCZ'])
        }
        print(f"🧬 GWAS ({row['ID_SCZ']}): Risk Allele (A1)=[{row['A1_SCZ']}], Other (A2)=[{row['A2_SCZ']}], Disease Beta=[{row['BETA_SCZ']}]")

    print("\n--- Scanning Raw EBI Tabix Data ---\n")
    
    # 2. Scan EBI Data
    with open(EBI_FILE, 'r') as f:
        next(f)
        for line in f:
            fields = line.rstrip('\n').split('\t')
            if len(fields) < 18:
                continue
                
            rsid = fields[-1]
            if rsid in TARGET_SNPS:
                
                # Identify Gene
                gene = "UNKNOWN"
                if "ENSG00000196917" in line: gene = "HCAR1"
                elif "ENSG00000182782" in line: gene = "HCAR2"
                elif "ENSG00000090975" in line: gene = "PITPNM2"
                
                if gene == "UNKNOWN": continue
                
                ebi_ref = str(fields[5]).upper()
                ebi_alt = str(fields[6]).upper()
                ebi_beta = float(fields[11])
                p_val = float(fields[10])
                
                # Only look at significant hits to avoid noise
                if p_val > 0.05: continue
                
                print(f"🔬 EBI ({rsid} -> {gene}): REF=[{ebi_ref}], ALT=[{ebi_alt}], EBI Beta (Effect of ALT)=[{ebi_beta}]")
                
                # Execute Alignment Logic
                gwas = gwas_dict[rsid]
                aligned_beta = None
                
                if gwas['A1_Risk'] == ebi_alt and gwas['A2_Ref'] == ebi_ref:
                    print(f"   ✅ Perfect Match: GWAS Risk ({gwas['A1_Risk']}) == EBI ALT ({ebi_alt}). Using EBI Beta.")
                    aligned_beta = ebi_beta
                elif gwas['A1_Risk'] == ebi_ref and gwas['A2_Ref'] == ebi_alt:
                    print(f"   🔄 Flipped Match: GWAS Risk ({gwas['A1_Risk']}) == EBI REF ({ebi_ref}). Multiplying EBI Beta by -1.")
                    aligned_beta = -ebi_beta
                else:
                    print(f"   ❌ Allele Mismatch/Strand Flip: GWAS[{gwas['A1_Risk']}/{gwas['A2_Ref']}] vs EBI[{ebi_ref}/{ebi_alt}]")
                    continue
                
                # Calculate Final Consequence
                print(f"   🧮 Math: Disease Beta ({gwas['SCZ_Beta']}) * Aligned eQTL Beta ({aligned_beta})")
                if (gwas['SCZ_Beta'] > 0 and aligned_beta > 0) or (gwas['SCZ_Beta'] < 0 and aligned_beta < 0):
                    print("   📈 Verdict: UP-REGULATES\n")
                else:
                    print("   📉 Verdict: COLLAPSES (Down-Regulates)\n")

if __name__ == "__main__":
    audit_directionality()
```
