## Supplemental Dataset 3 for "State-Dependent 3D Enhancer Architecture Resolves a Shared Schizophrenia and Multiple Sclerosis Ketone and Lactate Sensing Logic Gate": macrophage_pleiotropy_matcher-v2.py

```
import pandas as pd
import numpy as np

# Input Files
EBI_DUMP_FILE = "EBI_Immune_eQTL_Targeted_DeepSweep_Hits.tsv"
PLEIOTROPY_FILE = "SCZ_vs_IMSGC_MS_Pleiotropy_Analysis_Widened.tsv"
OUTPUT_FILE = "Macrophage_SCZ_MS_Smoking_Gun.tsv"

# The Ensembl IDs for the War
TARGET_GENES = {
    "ENSG00000196917": "HCAR1",   # Your Lactate Sensor
    "ENSG00000182782": "HCAR2",   # Your BHB Sensor
    "ENSG00000090975": "PITPNM2"  # The Reviewer's Pet Gene
}

# The Reviewer's Pet SNP
PET_SNP = "rs7975763"

def execute_macrophage_cross_reference():
    print("--- Initiating Robust Macrophage Pleiotropy Matcher ---")
    
    # 1. Load the Disease Pleiotropy Architecture
    print(f"Loading disease architecture: {PLEIOTROPY_FILE}...")
    try:
        df_pleio = pd.read_csv(PLEIOTROPY_FILE, sep='\t')
    except FileNotFoundError:
        print("❌ Error: Widened pleiotropy file not found.")
        return
        
    # Index for O(1) lookups
    df_pleio_sig = df_pleio[df_pleio['PVAL_SCZ'] < 1e-5]
    pleio_dict = df_pleio_sig.set_index('ID_SCZ').to_dict('index')
    print(f"✅ Indexed {len(pleio_dict)} highly significant shared MS/SCZ risk variants.")

    # 2. Parse the Raw Tabix Output (Robust Parsing)
    print(f"\nSifting through raw EBI database rows in {EBI_DUMP_FILE}...")
    
    matched_eqtls = []
    
    with open(EBI_DUMP_FILE, 'r') as f:
        next(f) # Skip header
        
        for line in f:
            fields = line.rstrip('\n').split('\t')
            
            # The absolute minimum columns EBI provides is 18 (plus our 2 prepended columns)
            if len(fields) < 18:
                continue
                
            # ROBUST GENE MATCH: Scan the entire line for the Ensembl ID
            # This bypasses EBI's habit of moving the Gene ID column around
            target_gene_name = None
            for ensg, name in TARGET_GENES.items():
                if ensg in line:
                    target_gene_name = name
                    break
                    
            if not target_gene_name:
                continue
                
            # ROBUST P-VALUE: Always index 10 (from the left)
            try:
                p_value = float(fields[10])
            except ValueError:
                continue
                
            if p_value > 0.05:
                continue
                
            # ROBUST RSID: Always the absolute last column (from the right)
            rsid = fields[-1]
            
            if rsid in pleio_dict:
                study = fields[0]
                disease_data = pleio_dict[rsid]
                
                # Extract EBI Alleles and Beta (Always fixed from the left)
                ebi_ref = str(fields[5]).upper()
                ebi_alt = str(fields[6]).upper()
                ebi_beta = float(fields[11])
                
                dis_a1 = str(disease_data['A1_SCZ']).upper()
                dis_a2 = str(disease_data['A2_SCZ']).upper()
                dis_beta = float(disease_data['BETA_SCZ'])
                
                # Align Beta
                if dis_a1 == ebi_alt and dis_a2 == ebi_ref:
                    aligned_eqtl_beta = ebi_beta
                elif dis_a1 == ebi_ref and dis_a2 == ebi_alt:
                    aligned_eqtl_beta = -ebi_beta
                else:
                    continue 
                
                if (dis_beta > 0 and aligned_eqtl_beta > 0) or (dis_beta < 0 and aligned_eqtl_beta < 0):
                    consequence = "UP-Regulates"
                else:
                    consequence = "COLLAPSES (Down-Regulates)"
                
                matched_eqtls.append({
                    'rsID': rsid,
                    'Target_Gene': target_gene_name,
                    'Macrophage_Study': study,
                    'Disease_PVAL': disease_data['PVAL_SCZ'],
                    'Regulatory_Region': disease_data['Regulatory_Region'],
                    'Macrophage_eQTL_PVAL': p_value,
                    'Biological_Consequence': consequence
                })

    if not matched_eqtls:
        print("\n⚠️ Zero overlaps found between disease risk and macrophage eQTLs.")
        return
        
    final_df = pd.DataFrame(matched_eqtls)
    final_df = final_df.sort_values(by='Disease_PVAL')
    
    print("\n=======================================================")
    print("🏆 THE VERDICT: MACROPHAGE TRANSCRIPTOMIC CROSS-REFERENCE")
    print("=======================================================")
    
    hcar1_hits = len(final_df[final_df['Target_Gene'] == 'HCAR1'])
    pitpnm2_hits = len(final_df[final_df['Target_Gene'] == 'PITPNM2'])
    
    print(f"\nTotal MS/SCZ Risk SNPs driving HCAR1 expression in Macrophages: {hcar1_hits}")
    print(f"Total MS/SCZ Risk SNPs driving PITPNM2 expression in Macrophages: {pitpnm2_hits}")
    
    if PET_SNP in final_df['rsID'].values:
        pet_targets = final_df[final_df['rsID'] == PET_SNP]['Target_Gene'].unique()
        print(f"\n🚨 ALERT: The Reviewer's Pet SNP ({PET_SNP}) was found!")
        print(f"In activated macrophages, it exclusively targets: {', '.join(pet_targets)}")
    
    print("\n--- TOP BIOLOGICAL SMOKING GUNS ---")
    print(final_df[['rsID', 'Target_Gene', 'Regulatory_Region', 'Macrophage_eQTL_PVAL', 'Biological_Consequence']].head(15).to_string(index=False))
    
    final_df.to_csv(OUTPUT_FILE, sep='\t', index=False)
    print(f"\n📁 Saved robust mapping to {OUTPUT_FILE}")

if __name__ == "__main__":
    execute_macrophage_cross_reference()
```
