## Supplemental Dataset 3 for "State-Dependent 3D Enhancer Architecture Resolves a Shared Schizophrenia and Multiple Sclerosis Ketone and Lactate Sensing Logic Gate": macrophage_pleiotropy_matcher.py

```
import pandas as pd
import numpy as np
import os

# Input Files
EBI_DUMP_FILE = "EBI_Immune_eQTL_Targeted_DeepSweep_Hits.tsv"
PLEIOTROPY_FILE = "SCZ_vs_IMSGC_MS_Pleiotropy_Analysis.tsv"

def execute_macrophage_cross_reference():
    print("--- Initiating Macrophage vs. Autoimmune/SCZ Cross-Reference ---")
    
    if not os.path.exists(EBI_DUMP_FILE) or not os.path.exists(PLEIOTROPY_FILE):
        print("⚠️ Error: Missing input files. Ensure both TSVs are in the directory.")
        return

    # 1. Load the data
    df_ebi = pd.read_csv(EBI_DUMP_FILE, sep='\t')
    df_pleio = pd.read_csv(PLEIOTROPY_FILE, sep='\t')
    
    # 2. Merge on the rsID
    print("Merging Macrophage eQTLs with the SCZ/MS Pleiotropy Landscape...")
    merged_df = pd.merge(
        df_ebi, 
        df_pleio, 
        left_on='rsID_Found', 
        right_on='ID_SCZ', 
        how='inner'
    )
    
    if merged_df.empty:
        print("\n❌ RESULT: None of the shared SCZ/MS variants appear in these macrophage datasets.")
        return
        
    # 3. APPLY STRICT SIGNIFICANCE FILTERS
    # Must be Genome-Wide Significant in SCZ and highly significant shared variance in MS
    print("Filtering for Genome-Wide Significant SCZ/MS drivers...")
    filtered_df = merged_df[
        (merged_df['PVAL_SCZ'] < 5e-8) & 
        (merged_df['PVAL_MS'] < 1e-4)
    ].copy()
    
    if filtered_df.empty:
        print("\n❌ RESULT: We found overlaps, but none of the strictly significant SCZ/MS peaks hit the macrophages.")
        return

    # 4. CALCULATE THE DIRECTION OF TRANSCRIPTION
    # By multiplying the Disease Beta by the Macrophage Beta, we align the effect to the Risk Allele.
    directions = []
    for _, row in filtered_df.iterrows():
        scz_beta = float(row['BETA_SCZ'])
        macro_beta = float(row['Beta (Effect)'])
        
        # If signs are opposite (Product < 0), the Risk Allele suppresses the gene.
        if (scz_beta * macro_beta) < 0:
            directions.append("DOWN-Regulated (Switch Broken)")
        else:
            directions.append("UP-Regulated (Switch Stuck On)")
            
    filtered_df['Transcriptomic_Consequence'] = directions

    # 5. Clean up the output to tell a clear biological story
    output_cols = [
        'Regulatory_Region',
        'Gene',
        'rsID_Found',
        'Cell_Type',
        'Condition',
        'P_Value',          # Macrophage eQTL P-value
        'Beta (Effect)',    # Macrophage Beta
        'Transcriptomic_Consequence',
        'PVAL_SCZ',
        'PVAL_MS',
        'Pleiotropic_Relationship'
    ]
    
    final_df = filtered_df[output_cols].copy()
    final_df = final_df.rename(columns={
        'P_Value': 'PVAL_Macrophage_eQTL',
        'Beta (Effect)': 'BETA_Macrophage_eQTL'
    })
    
    # Sort by the most significant Macrophage disruption
    # Drop duplicates just in case multiple datasets recorded the exact same event
    final_df = final_df.sort_values(by=['Gene', 'PVAL_Macrophage_eQTL'])
    final_df = final_df.drop_duplicates(subset=['rsID_Found', 'Gene', 'Cell_Type', 'Condition'])
    
    # 6. Export the final smoking gun
    out_file = "Macrophage_SCZ_MS_Smoking_Gun.tsv"
    final_df.to_csv(out_file, sep='\t', index=False)
    
    print(f"\n✅ CROSS-REFERENCE SUCCESS! Found {len(final_df)} overlapping biological events.")
    print(f"📁 Results saved to: {out_file}")
    
    print("\n--- THE SMOKING GUN: TRUE SCZ/MS DRIVERS IN MACROPHAGES ---")
    preview_cols = ['Regulatory_Region', 'Gene', 'Cell_Type', 'Condition', 'Transcriptomic_Consequence', 'PVAL_Macrophage_eQTL']
    print(final_df[preview_cols].head(15).to_string(index=False))

if __name__ == "__main__":
    execute_macrophage_cross_reference()
```
