## Supplemental Dataset 3 for "State-Dependent 3D Enhancer Architecture Resolves a Shared Schizophrenia and Multiple Sclerosis Ketone and Lactate Sensing Logic Gate": macrophage_state_decoder.py

```
import pandas as pd

MANIFEST_FILE = "tabix_ftp_paths.tsv"
SMOKING_GUN_FILE = "Macrophage_SCZ_MS_Smoking_Gun.tsv"
OUTPUT_FILE = "Macrophage_SCZ_MS_Smoking_Gun_Decoded.tsv"

def decode_states():
    print("--- Initiating EBI QTS/QTD Metadata Decoder ---\n")
    
    # 1. Load the EBI Master Manifest
    try:
        df_manifest = pd.read_csv(MANIFEST_FILE, sep='\t')
    except FileNotFoundError:
        print(f"❌ Error: Could not find {MANIFEST_FILE}")
        return
        
    # Dynamically find the right columns in the manifest
    cols = [c.lower() for c in df_manifest.columns]
    
    try:
        study_col = df_manifest.columns[['study' in c for c in cols].index(True)]
        
        # 'qtl_group' or 'condition_label' usually holds the activation state (naive, LPS, etc)
        if 'condition_label' in cols:
            cond_col = df_manifest.columns[cols.index('condition_label')]
        elif 'qtl_group' in cols:
            cond_col = df_manifest.columns[cols.index('qtl_group')]
        else:
            cond_col = df_manifest.columns[2] # Fallback
            
        # 'tissue_label' usually holds monocyte/macrophage
        if 'tissue_label' in cols:
            cell_col = df_manifest.columns[cols.index('tissue_label')]
        else:
            cell_col = df_manifest.columns[3] # Fallback
            
    except ValueError:
        print("❌ Error: Manifest column headers are non-standard.")
        return

    # Build a fast dictionary mapping QTS ID -> "Macrophage | Salmonella"
    # Note: Some QTS studies have multiple QTD datasets, so we map to the exact FTP path if possible, 
    # but the simple study ID is a good fallback for the overarching cohort.
    print("Mapping metadata dictionary...")
    meta_dict = {}
    for _, row in df_manifest.iterrows():
        study_id = str(row[study_col]).strip()
        cell_type = str(row[cell_col]).strip()
        condition = str(row[cond_col]).strip()
        meta_dict[study_id] = f"{cell_type} | {condition}"

    # 2. Load the Smoking Gun Data
    print("Loading Smoking Gun eQTLs...")
    df_hits = pd.read_csv(SMOKING_GUN_FILE, sep='\t')

    # 3. Apply the Decoding
    decoded_states = []
    for qts in df_hits['Macrophage_Study']:
        decoded_states.append(meta_dict.get(qts, "Unknown State"))
        
    df_hits['Biological_State'] = decoded_states
    
    # Reorder columns for readability, keeping insightful ones up front but RETAINING ALL RAW DATA
    front_cols = ['rsID', 'Target_Gene', 'Biological_State', 'Biological_Consequence', 'Macrophage_eQTL_PVAL', 'Regulatory_Region']
    remaining_cols = [c for c in df_hits.columns if c not in front_cols]
    df_hits = df_hits[front_cols + remaining_cols]
    
    # Save the full decoded file
    df_hits.to_csv(OUTPUT_FILE, sep='\t', index=False)
    print(f"✅ Full decoded dataset saved to: {OUTPUT_FILE}\n")
 
    # 4. Generate the Biological Summary Pivot
    print("=======================================================")
    print("🧬 STATE-DEPENDENT TRANSCRIPTOMIC SUMMARY")
    print("=======================================================")
    
    # Group by Gene, State, and Consequence to see the patterns
    summary = df_hits.groupby(['Target_Gene', 'Biological_State', 'Biological_Consequence']).size().reset_index(name='Hit_Count')
    
    # Sort for easy reading
    summary = summary.sort_values(by=['Target_Gene', 'Biological_State'])
    print(summary.to_string(index=False))
    
    print("\n=======================================================")
    
if __name__ == "__main__":
    decode_states()
```
