## Supplemental Dataset 3 for "State-Dependent 3D Enhancer Architecture Resolves a Shared Schizophrenia and Multiple Sclerosis Ketone and Lactate Sensing Logic Gate": README.docx

**Supplemental Dataset 3: High-Resolution Immune eQTL Transcriptomic Fine-Mapping**

**Manuscript Title:** State-Dependent 3D Enhancer Architecture Resolves a Shared Schizophrenia and Multiple Sclerosis Ketone and Lactate Sensing Logic Gate

**Principal Investigator:** Bryan A. Krantz, Ph.D.

**Laboratory:** The Krantz Lab, University of Maryland, Baltimore

**Overview**

This directory contains the computational pipeline and raw datasets used to definitively map the 3D regulatory architecture of the *HCAR* locus (Chromosome 12: 122.80 - 123.70 Mb).

To overcome legacy 1D proximity heuristics (e.g., FUMA) and bypass server-side artifacts/throttling inherent to standard REST API queries, this pipeline utilizes remote byte-range **Tabix extractions** directly against the EMBL-EBI eQTL Catalogue FTP servers. This rigorous approach ensures absolute strand-alignment integrity and allows us to isolate macrophage-specific transcriptomic responses to distinct pathogen-associated molecular patterns (PAMPs).

**Pipeline Execution Order**

**1. Data Extraction (The Tabix Bypass)**

- **remote_ftp_tabix_miner_hg38_liftover.py**: The primary extraction engine. Bypasses the EBI REST API to stream raw eQTL data directly from EBI FTP servers. It targets the 3D structural domain (122.80–123.70 Mb, converted to hg38 liftover coordinates) across isolated macrophage and monocyte lineages.
- **tabix_ftp_paths.tsv**: The master manifest of EBI FTP endpoints and study metadata used by the miner.
- *Output:* EBI_Immune_eQTL_Targeted_DeepSweep_Hits.tsv (Raw, unfiltered genomic extraction).

**2. Disease Risk Cross-Referencing**

- **SCZ_vs_IMSGC_MS_Pleiotropy_Analysis_Widened.tsv**: The baseline disease architecture containing the highly significant, shared risk variants for Schizophrenia (PGC3) and Multiple Sclerosis (IMSGC) across the widened topological domain.
- **macrophage_pleiotropy_matcher_v3.py**: Cross-references the raw EBI eQTL dump against our shared disease risk alleles. Crucially, this script rigorously aligns the eQTL Beta (effect size) to the established SCZ/MS risk strand to determine true biological directionality (Up-regulation vs. Collapse).
- *Output:* Macrophage_SCZ_MS_Smoking_Gun.tsv

**3. Biological State Decoding**

- **macrophage_state_decoder.py**: Maps the cryptic EBI study IDs (QTS identifiers) to specific, experimentally controlled biological states (e.g., Naive, *Salmonella* LPS, MDP, IFN-γ).
- *Output:* Macrophage_SCZ_MS_Smoking_Gun_Decoded.tsv (The final, human-readable master dataset).

**4. Architectural and Thermodynamic Auditing**

- **architectural_enhancer_audit.py**: Analyzes the decoded dataset to verify the spatial origin of the eQTLs, confirming that the 123.60 Mb distal hub physically targets the *HCAR* array and exerts zero significant regulatory control over the adjacent *PITPNM2* structural gene.
- **audit_directionality.py**: A statistical diagnostic script that confirms the state-dependent logic gate: under Gram-negative stress, the disease risk variants collapses *HCAR2* and *HCAR1*.

**Key Data Files**

- **Macrophage_SCZ_MS_Smoking_Gun_Decoded.tsv**: The definitive dataset supporting Figure 3 and Figure 4 of the manuscript. It contains the exact rsIDs, target genes, pathogen states, and aligned transcriptional consequences proving the breakdown of the thermodynamic logic gate.

*(Note: Deprecated REST API scripts such as ebi_eqtl_macrophage_miner.py have been retained in this directory solely for historical version control and demonstration of their brittle interface.)*
