## Supplemental Dataset 3 for "State-Dependent 3D Enhancer Architecture Resolves a Shared Schizophrenia and Multiple Sclerosis Ketone and Lactate Sensing Logic Gate": remote_ftp_tabix_miner.py

```
import pandas as pd
import subprocess
import time
import sys

MANIFEST_FILE = "tabix_ftp_paths.tsv"
OUTPUT_FILE = "EBI_Immune_eQTL_Targeted_DeepSweep_Hits.tsv"

# The expanded physical TAD window (Captures HCAR1/2 and PITPNM2 simultaneously)
REGION = "12:122800000-123700000"

def run_remote_tabix():
    print("--- Initiating EMBL-EBI eQTL Macrophage Miner (REMOTE FTP TABIX OVERRIDE) ---")
    
    # 1. Load the manifest
    print(f"Loading master manifest: {MANIFEST_FILE}")
    try:
        df = pd.read_csv(MANIFEST_FILE, sep='\t')
    except FileNotFoundError:
        print(f"❌ Error: {MANIFEST_FILE} not found. Ensure you are in the Supplemental_Dataset_3 directory.")
        sys.exit(1)
        
    # 2. Find Macrophage / Monocyte tracks dynamically
    mask = df.astype(str).apply(lambda x: x.str.contains('macrophage|monocyte', case=False)).any(axis=1)
    immune_df = df[mask]
    
    print(f"✅ Isolated {len(immune_df)} specific Macrophage/Monocyte datasets.")
    print(f"📡 Bypassing REST API. Streaming remote byte-ranges for {REGION} directly from EBI FTP servers...\n")
    
    # Identify the FTP path and Study columns dynamically based on standard EBI headers
    try:
        ftp_col = [col for col in df.columns if 'ftp' in col.lower() or 'url' in col.lower()][0]
        study_col = [col for col in df.columns if 'study' in col.lower()][0]
    except IndexError:
        print("❌ Error: Could not identify 'ftp' or 'study' columns in the manifest.")
        sys.exit(1)
    
    total_hits = 0
    
    with open(OUTPUT_FILE, 'w') as out_f:
        # Write a generic header (Tabix pulls raw VCF/TSV lines without headers)
        out_f.write("STUDY\tFTP_PATH\tRAW_TABIX_OUTPUT\n")
        
        for index, row in immune_df.iterrows():
            ftp_url = row[ftp_col]
            study_name = row[study_col]
            
            print(f"-> Streaming Tabix: {study_name} | {ftp_url.split('/')[-1]}...")
            
            # The tabix command: tabix ftp://.../file.sumstats.gz 12:122800000-123700000
            cmd = ["tabix", ftp_url, REGION]
            
            try:
                # Run the command and capture the output
                result = subprocess.run(cmd, capture_output=True, text=True, check=True)
                
                hits = result.stdout.strip().split('\n')
                hits = [h for h in hits if h] # Remove empty strings
                
                if hits:
                    print(f"   🔥 FOUND {len(hits)} eQTL variants in this 3D block!")
                    total_hits += len(hits)
                    for hit in hits:
                        out_f.write(f"{study_name}\t{ftp_url}\t{hit}\n")
                else:
                    print(f"   - Zero hits in this window.")
                    
            except subprocess.CalledProcessError:
                print(f"   ⚠️ Tabix stream failed for this dataset (FTP timeout or bad index).")
            except FileNotFoundError:
                print("❌ CRITICAL ERROR: 'tabix' command not found on your system.")
                print("   Install it via Homebrew: `brew install htslib`")
                sys.exit(1)
            
            # Brief 0.5s sleep to avoid triggering FTP DDoS protections
            time.sleep(0.5)

    print(f"\n=======================================================")
    print(f"✅ REMOTE SWEEP COMPLETE.")
    print(f"🎯 Total eQTL hits extracted across all datasets: {total_hits}")
    print(f"📁 Raw data saved to: {OUTPUT_FILE}")
    print(f"Note: Because Tabix pulls raw database rows instead of JSON, you may need to adjust")
    print(f"the column indexing in macrophage_pleiotropy_matcher.py to target the correct P-value/Beta columns.")
    print(f"=======================================================")

if __name__ == "__main__":
    run_remote_tabix()
```
