## Supplemental Dataset 4 for "State-Dependent 3D Enhancer Architecture Resolves a Shared Schizophrenia and Multiple Sclerosis Ketone and Lactate Sensing Logic Gate": ld_block_distance_calculator.py

```
import pandas as pd
import os

def calculate_ld_distance():
    print("--- Calculating 3D LD Block Physical Distance ---")
    
    rsid_file = "LDLink_Target_rsIDs.txt"
    tsv_file = "SCZ_vs_IMSGC_MS_Pleiotropy_Analysis_Widened.tsv"
    
    if not os.path.exists(rsid_file):
        print(f"❌ Error: Could not find '{rsid_file}'.")
        return
    if not os.path.exists(tsv_file):
        print(f"❌ Error: Could not find '{tsv_file}'.")
        return

    with open(rsid_file, 'r') as f:
        target_rsids = [line.strip() for line in f if line.strip()]
        
    print(f"Loaded {len(target_rsids)} target rsIDs from the linkage set.")
    
    print(f"Scanning '{tsv_file}' for coordinate mappings...")
    df = pd.read_csv(tsv_file, sep='\t', low_memory=False)
    
    # Intelligently find the ID and Position columns based on your typical pipeline naming
    id_col = next((c for c in df.columns if c.upper() in ['ID', 'SNP', 'RSID', 'ID_SCZ', 'MARKERNAME']), None)
    pos_col = next((c for c in df.columns if c.upper() in ['POS', 'BP', 'POSITION', 'BP_SCZ']), None)
    
    if not id_col or not pos_col:
        print("❌ Error: Could not confidently identify the rsID or Base Pair Position columns.")
        print("Available columns:", list(df.columns))
        return

    matched_df = df[df[id_col].isin(target_rsids)].copy()
    
    if matched_df.empty:
        print("❌ Error: None of the target rsIDs were found in the TSV.")
        return
        
    # Force numeric conversion just in case there are string artifacts
    matched_df[pos_col] = pd.to_numeric(matched_df[pos_col], errors='coerce')
    matched_df = matched_df.dropna(subset=[pos_col])
    
    # Identify the boundaries
    min_bp = int(matched_df[pos_col].min())
    max_bp = int(matched_df[pos_col].max())
    distance_bp = max_bp - min_bp
    distance_kb = distance_bp / 1000.0
    
    # Identify the specific boundary SNPs
    min_snp = matched_df[matched_df[pos_col] == min_bp][id_col].values[0]
    max_snp = matched_df[matched_df[pos_col] == max_bp][id_col].values[0]

    print("\n==================================================")
    print("🧬 LD BLOCK ARCHITECTURE METRICS")
    print("==================================================")
    print(f"Target SNPs Matched : {len(matched_df)} / {len(target_rsids)}")
    print(f"5' Boundary (Min)   : {min_bp:,} ({min_snp})")
    print(f"3' Boundary (Max)   : {max_bp:,} ({max_snp})")
    print("--------------------------------------------------")
    print(f"Total Block Distance: {distance_bp:,} base pairs")
    print(f"                      {distance_kb:,.2f} kb")
    print("==================================================\n")
    
    if distance_kb > 450 and distance_kb < 550:
        print("✅ Validation: This confirms the '~500kb' linkage block text in the manuscript is mathematically perfect!")

if __name__ == "__main__":
    calculate_ld_distance()
```
