## Supplemental Dataset 4 for "State-Dependent 3D Enhancer Architecture Resolves a Shared Schizophrenia and Multiple Sclerosis Ketone and Lactate Sensing Logic Gate": LDLink_rsID_miner.py

```
import pandas as pd

# Input file
PLEIO_FILE = "SCZ_vs_IMSGC_MS_Pleiotropy_Analysis_Widened.tsv"
OUTPUT_FILE = "LDLink_Target_rsIDs.txt"

# The Reviewer's Pet SNP that must be included
PET_SNP = "rs7975763"

def extract_ldlink_targets():
    print("--- Initiating High-Value rsID Miner for LDLink ---\n")
    
    try:
        df = pd.read_csv(PLEIO_FILE, sep='\t')
    except FileNotFoundError:
        print(f"❌ Error: {PLEIO_FILE} not found.")
        return

    # Ensure we only look at variants with high significance
    df_sig = df[df['PVAL_SCZ'] < 1e-5].copy()
    
    # Sort the entire dataframe by SCZ P-value (strongest hits at the top)
    df_sig = df_sig.sort_values(by='PVAL_SCZ', ascending=True)

    regions = [
        "5' Skyscraper (HCAR2)",
        "3' Skyscraper (HCAR1)",
        "Distal Enhancer (PITPNM2 / Reviewer's Peak)"
    ]

    target_rsids = []
    
    # 1. Force the Reviewer's Pet SNP to the very top of the list
    if PET_SNP in df_sig['ID_SCZ'].values:
        target_rsids.append(PET_SNP)
        print(f"📌 Pinned Reviewer's Pet SNP: {PET_SNP}")
    else:
        print(f"⚠️ Warning: {PET_SNP} not found in the significant dataset!")

    print("\nExtracting Top 5 Index SNPs per Regulatory Region:")
    
    # 2. Loop through each region and grab the top 5 strongest statistical tags
    for region in regions:
        region_df = df_sig[df_sig['Regulatory_Region'] == region]
        
        # Get top 5, excluding the Pet SNP if it was already grabbed
        top_snps = region_df[~region_df['ID_SCZ'].isin(target_rsids)].head(5)
        
        print(f"\n📍 Region: {region}")
        for _, row in top_snps.iterrows():
            rsid = row['ID_SCZ']
            pval = row['PVAL_SCZ']
            target_rsids.append(rsid)
            print(f"   -> {rsid} (SCZ P-val: {pval:.2e})")

    # 3. Export to a clean text file for easy copy-pasting into the NIH LDLink Website
    with open(OUTPUT_FILE, 'w') as f:
        for rsid in target_rsids:
            f.write(f"{rsid}\n")
            
    print(f"\n✅ Successfully exported {len(target_rsids)} high-value rsIDs to {OUTPUT_FILE}")
    print("📋 You can copy and paste the contents of this file directly into the LDMatrix tool.")

if __name__ == "__main__":
    extract_ldlink_targets()
```
