## Supplemental Dataset 4 for "State-Dependent 3D Enhancer Architecture Resolves a Shared Schizophrenia and Multiple Sclerosis Ketone and Lactate Sensing Logic Gate": LDLink_rsID_to_Test.rtf

(tf-macos) bakrantz@Bryans-MacBook-Pro Supplemental_Dataset_4 % python LDLink_rsID_miner.py 
--- Initiating High-Value rsID Miner for LDLink ---

?? Pinned Reviewer's Pet SNP: rs7975763

Extracting Top 5 Index SNPs per Regulatory Region:

?? Region: 5' Skyscraper (HCAR2)
   -> rs4758662 (SCZ P-val: 2.53e-07)
   -> rs4758643 (SCZ P-val: 2.68e-07)
   -> rs34129612 (SCZ P-val: 2.82e-07)
   -> rs7135419 (SCZ P-val: 2.87e-07)
   -> rs10773152 (SCZ P-val: 3.02e-07)

?? Region: 3' Skyscraper (HCAR1)
   -> rs3741530 (SCZ P-val: 2.78e-19)
   -> rs55742290 (SCZ P-val: 3.38e-19)
   -> rs28755851 (SCZ P-val: 4.16e-19)
   -> rs883562 (SCZ P-val: 4.37e-19)
   -> rs3759115 (SCZ P-val: 4.37e-19)

?? Region: Distal Enhancer (PITPNM2 / Reviewer's Peak)
   -> rs1790135 (SCZ P-val: 6.91e-24)
   -> rs2102949 (SCZ P-val: 7.05e-24)
   -> rs4460848 (SCZ P-val: 9.12e-24)
   -> rs1716180 (SCZ P-val: 9.23e-24)
   -> rs2851447 (SCZ P-val: 1.03e-23)

✅ Successfully exported 16 high-value rsIDs to LDLink_Target_rsIDs.txt
?? You can copy and paste the contents of this file directly into the LDMatrix tool.

Populations (CEU) (TSI) (FIN) (GBR) (IBS)

LDMatrix refs

Machiela MJ, Chanock SJ. LDlink: a web-based application for exploring population-specific haplotype structure and linking correlated alleles of possible functional variants. Bioinformatics. 2015 Jul 2.

Breeze, C.E., Haugen, E., Gutierrez-Arcelus, M., Yao, X., Teschendorff, A., Beck, S., Dunham, I., Stamatoyannopoulos, J., Franceschini, N., Machiela, M.J., Berndt, S.I. FORGEdb: a tool for identifying candidate functional variants and uncovering target genes and mechanisms for complex diseases. Genome Biology. 25, 3 (2024).
