## Supplemental Dataset 5 for "State-Dependent 3D Enhancer Architecture Resolves a Shared Schizophrenia and Multiple Sclerosis Ketone and Lactate Sensing Logic Gate": coloc_output.txt

(tf-macos) bakrantz@Bryans-MacBook-Pro Supplemental_Dataset_5 % Rscript run_coloc_analysis.R 

=======================================================
🧬 BAYESIAN COLOCALIZATION ANALYSIS
=======================================================

--- Executing COLOC for HCAR1 (315 SNPs) ---
📊 Posterior Probabilities:
  PP.H3 (Distinct Variants / LD Bleed) : 0.0744
  PP.H4 (Shared Causal Variant)        : 0.4509
⚠️ VERDICT: INCONCLUSIVE (Neither H3 nor H4 > 0.8).

--- Executing COLOC for HCAR2 (415 SNPs) ---
📊 Posterior Probabilities:
  PP.H3 (Distinct Variants / LD Bleed) : 0.1365
  PP.H4 (Shared Causal Variant)        : 0.2756
⚠️ VERDICT: INCONCLUSIVE (Neither H3 nor H4 > 0.8).

--- Executing COLOC for PITPNM2 (250 SNPs) ---
📊 Posterior Probabilities:
  PP.H3 (Distinct Variants / LD Bleed) : 0.0569
  PP.H4 (Shared Causal Variant)        : 0.2913
⚠️ VERDICT: INCONCLUSIVE (Neither H3 nor H4 > 0.8).
Warning messages:
1: In check_dataset(d = dataset2, 2) : minimum p value is: 0.0014515
If this is what you expected, this is not a problem.
If this is not as small as you expected, please check you supplied var(beta) and not sd(beta) for the varbeta argument. If that's not the explanation, please check the 02_data vignette.
2: In check_dataset(d = dataset2, 2) : minimum p value is: 0.0005981
If this is what you expected, this is not a problem.
If this is not as small as you expected, please check you supplied var(beta) and not sd(beta) for the varbeta argument. If that's not the explanation, please check the 02_data vignette.
3: In check_dataset(d = dataset2, 2) : minimum p value is: 0.00045882
If this is what you expected, this is not a problem.
If this is not as small as you expected, please check you supplied var(beta) and not sd(beta) for the varbeta argument. If that's not the explanation, please check the 02_data vignette.

=======================================================
Export complete. Use PP.H4 values for the manuscript resubmission.
