## Supplemental Dataset 5 for "State-Dependent 3D Enhancer Architecture Resolves a Shared Schizophrenia and Multiple Sclerosis Ketone and Lactate Sensing Logic Gate": colocalization_data_prepper.py

```
import pandas as pd
import os

# Input Files
GWAS_FILE = "SCZ_vs_IMSGC_MS_Pleiotropy_Analysis_Widened.tsv"
EQTL_FILE = "Macrophage_SCZ_MS_Smoking_Gun_Decoded.tsv"
OUTPUT_FILE = "coloc_input_HCAR1.tsv"

# Standard Sample Sizes (N)
# PGC3 Schizophrenia Wave 3 is approx 130,000. 
# Average EBI Macrophage cohort (e.g. BLUEPRINT/Nedelec) is approx 500.
N_GWAS_SCZ = 130000 
N_EQTL_MACROPHAGE = 500

def prep_coloc_data():
    print("--- Initiating Bayesian Colocalization (COLOC) Prepper ---")
    
    if not os.path.exists(GWAS_FILE) or not os.path.exists(EQTL_FILE):
        print("⚠️ Error: Missing input files in the current directory.")
        return
        
    print("Loading GWAS and eQTL datasets...")
    df_gwas = pd.read_csv(GWAS_FILE, sep='\t')
    df_eqtl = pd.read_csv(EQTL_FILE, sep='\t')
    
    # Target genes to run separate COLOC analyses on
    targets = ['HCAR1', 'HCAR2', 'PITPNM2']
    
    for gene in targets:
        print(f"\n--- Processing {gene} ---")
        df_eqtl_gene = df_eqtl[df_eqtl['Target_Gene'] == gene].copy()
        
        if df_eqtl_gene.empty:
            print(f"⚠️ No significant eQTLs found for {gene}. COLOC cannot be run (PP.H4 effectively 0).")
            continue
            
        print(f"Isolated {len(df_eqtl_gene)} variants driving {gene} expression.")
        
        # Merge the datasets on rsID
        merged_df = pd.merge(
            df_gwas, 
            df_eqtl_gene, 
            left_on='ID_SCZ', 
            right_on='rsID', 
            how='inner'
        )
        
        if merged_df.empty:
            print(f"⚠️ No overlapping GWAS/eQTL variants for {gene}.")
            continue
            
        # Format the required columns for R's coloc.abf()
        coloc_df = pd.DataFrame()
        coloc_df['snp'] = merged_df['rsID']
        coloc_df['pval_gwas'] = merged_df['PVAL_SCZ']
        coloc_df['pval_eqtl'] = merged_df['Macrophage_eQTL_PVAL']
        coloc_df['N_gwas'] = N_GWAS_SCZ
        coloc_df['N_eqtl'] = N_EQTL_MACROPHAGE
        
        # Handle MAF
        if 'MAF' in merged_df.columns:
            coloc_df['MAF'] = merged_df['MAF']
        elif 'FREQ' in merged_df.columns:
            coloc_df['MAF'] = merged_df['FREQ']
        else:
            coloc_df['MAF'] = 0.10 # Safe placeholder for European GWAS
            
        # Remove duplicates
        coloc_df = coloc_df.drop_duplicates(subset=['snp'])
        
        output_name = f"coloc_input_{gene}.tsv"
        coloc_df.to_csv(output_name, sep='\t', index=False)
        print(f"✅ Prepped {len(coloc_df)} SNPs -> Saved to {output_name}")

    print("\nNext Step: Run these TSVs through the R coloc.abf() function.")

if __name__ == "__main__":
    prep_coloc_data()
```
