## Supplemental Dataset 5 for "State-Dependent 3D Enhancer Architecture Resolves a Shared Schizophrenia and Multiple Sclerosis Ketone and Lactate Sensing Logic Gate": README.docx

**Supplemental Dataset 5: Bayesian Colocalization (COLOC) Analysis**

**Manuscript Title:** State-Dependent 3D Enhancer Architecture Resolves a Shared Schizophrenia and Multiple Sclerosis Ketone and Lactate Sensing Logic Gate

**Principal Investigator:** Bryan A. Krantz, Ph.D.

**Laboratory:** The Krantz Lab, University of Maryland, Baltimore

**Overview**

This directory contains the computational scripts, input datasets, and output logs for the Bayesian colocalization analysis (coloc) requested during the peer review process.

The goal of this analysis was to test the hypothesis that the Schizophrenia/Multiple Sclerosis disease risk variants and the macrophage eQTL signals (for *HCAR1*, *HCAR2*, and *PITPNM2*) are driven by the exact same underlying causal variant (Hypothesis 4, PP.H4), rather than being independent signals in linkage disequilibrium (Hypothesis 3, PP.H3).

**Directory Contents**

**1. Execution Scripts**

- **colocalization_data_prepper.py**: Python script that merges the widened MS/SCZ pleiotropy architecture with the decoded EBI eQTL dataset. It generates standard input files formatted specifically for the R coloc package.
- **run_coloc_analysis.R**: The R script that loads the coloc library and calculates the Approximate Bayes Factors (ABF) for each of our target genes.

**2. Input Datasets**

- **coloc_input_HCAR1.tsv**: Formatted GWAS/eQTL overlap data for *HCAR1*.
- **coloc_input_HCAR2.tsv**: Formatted GWAS/eQTL overlap data for *HCAR2*.
- **coloc_input_PITPNM2.tsv**: Formatted GWAS/eQTL overlap data for *PITPNM2* (The reviewer's specified target).
- *Note: Source datasets (Macrophage_SCZ_MS_Smoking_Gun_Decoded.tsv and SCZ_vs_IMSGC_MS_Pleiotropy_Analysis_Widened.tsv) are provided here for reproducibility.*

**3. Results**

- **coloc_output.txt**: The raw console output and posterior probabilities generated by the R script.

**Interpretation of Results**

As detailed in coloc_output.txt, the Bayesian colocalization analysis yielded **inconclusive** posterior probabilities for a single shared causal variant (PP.H4 < 0.8) across all three tested genes (*HCAR1*, *HCAR2*, and *PITPNM2*). Furthermore, the probability of distinct, independent variants (PP.H3) was also extremely low (< 0.15).

**Biological Context and Limitations of COLOC:**

Standard Bayesian colocalization algorithms (coloc.abf) are mathematically optimized for simple 1D genomic architectures, assuming a single causal variant operating in a static cellular state.

However, as demonstrated in our primary manuscript, the 122.80–123.70 Mb locus does not operate as a single static variant. It functions as a massive, 772kb **3D distal pleiotropic enhancer hub** that dynamically shifts its physical looping based on the specific pathogen-associated molecular pattern (PAMP) (e.g., LPS vs. MDP).

Because the transcriptomic consequences of these variants are highly state-dependent (the effect between *HCAR1* and *HCAR2*), aggregating these dynamic states into a single Bayesian prior inherently dilutes the statistical power of the colocalization model.

Crucially, the failure of *PITPNM2* to achieve colocalization (PP.H4 = 0.29) further refutes the legacy 1D heuristic that assigned the 123.60 Mb distal hub exclusively to this structural gene. Ultimately, this underscores the necessity of relying on direct, state-dependent 3D transcriptomic mapping (Supplemental Dataset 3) rather than static 1D Bayesian approximations when evaluating complex immune logic gates.
