## Supplemental Dataset 5 for "State-Dependent 3D Enhancer Architecture Resolves a Shared Schizophrenia and Multiple Sclerosis Ketone and Lactate Sensing Logic Gate": run_coloc_analysis.R

```
suppressPackageStartupMessages(library(coloc))

targets <- c("HCAR1", "HCAR2", "PITPNM2")

run_coloc_for_gene <- function(gene) {
file_name <- paste0("coloc_input_", gene, ".tsv")
if (!file.exists(file_name)) {
    cat(sprintf("⚠️  Warning: %s not found. Skipping...\n", file_name))
    return(NULL)
}

# Load the prepped data
df <- read.table(file_name, header = TRUE, sep = "\t", stringsAsFactors = FALSE)

# Ensure there are no NA values in crucial columns (P-values and MAF)
df <- df[complete.cases(df[, c("pval_gwas", "pval_eqtl", "MAF")]), ]

if (nrow(df) == 0) {
    cat(sprintf("⚠️  Warning: No valid variants left for %s after NA removal.\n", gene))
    return(NULL)
}

cat(sprintf("\n--- Executing COLOC for %s (%d SNPs) ---\n", gene, nrow(df)))

# STREAMING_CHUNK:Formatting datasets for coloc.abf...

# Dataset 1: Schizophrenia GWAS
# type="cc" (case-control), s=0.3 (approx proportion of cases in PGC3)
dataset_gwas <- list(
    pvalues = df$pval_gwas,
    N = df$N_gwas[1],
    MAF = df$MAF,
    type = "cc",
    s = 0.3,
    snp = df$snp
)

# Dataset 2: Macrophage eQTL (EBI)
# type="quant" (quantitative trait for gene expression)
dataset_eqtl <- list(
    pvalues = df$pval_eqtl,
    N = df$N_eqtl[1],
    MAF = df$MAF,
    type = "quant",
    snp = df$snp
)

# STREAMING_CHUNK:Executing Approximate Bayes Factor calculation...

# Run the Bayesian Colocalization
# Note: Suppressing the standard print output to format it cleanly below
capture.output(
    my_coloc <- coloc.abf(dataset1 = dataset_gwas, dataset2 = dataset_eqtl)
)

# Extract the Posterior Probabilities
pp_h0 <- my_coloc$summary["PP.H0.abf"] # No association
pp_h1 <- my_coloc$summary["PP.H1.abf"] # GWAS only
coloc_h2 <- my_coloc$summary["PP.H2.abf"] # eQTL only
pp_h3 <- my_coloc$summary["PP.H3.abf"] # Two distinct causal variants (The Reviewer's assumption)
pp_h4 <- my_coloc$summary["PP.H4.abf"] # ONE shared causal variant (The Vanguard reality)

# STREAMING_CHUNK:Formatting and printing the verdict...

cat("📊 Posterior Probabilities:\n")
cat(sprintf("  PP.H3 (Distinct Variants / LD Bleed) : %.4f\n", pp_h3))
cat(sprintf("  PP.H4 (Shared Causal Variant)        : %.4f\n", pp_h4))

if (pp_h4 > 0.8) {
    cat(sprintf("🏆 VERDICT: STRONG COLOCALIZATION (PP.H4 > 0.8).\n"))
    cat(sprintf("   The disease risk and %s expression are driven by the SAME variant.\n", gene))
} else if (pp_h3 > 0.8) {
    cat(sprintf("❌ VERDICT: INDEPENDENT SIGNALS (PP.H3 > 0.8).\n"))
    cat(sprintf("   The disease peak and %s eQTL are distinct mutations in LD.\n", gene))
} else {
    cat(sprintf("⚠️ VERDICT: INCONCLUSIVE (Neither H3 nor H4 > 0.8).\n"))
}

}

cat("\n=======================================================\n")
cat("🧬 BAYESIAN COLOCALIZATION ANALYSIS")
cat("\n=======================================================\n")

for (g in targets) {
run_coloc_for_gene(g)
}

cat("\n=======================================================\n")
cat("Export complete. Use PP.H4 values for the manuscript resubmission.\n")
```
